# A genetically-encoded toolkit of functionalized nanobodies against fluorescent proteins for visualizing and manipulating intracellular signalling

**DOI:** 10.1101/544700

**Authors:** David L. Prole, Colin W. Taylor

## Abstract

**Background:** Intrabodies enable targeting of proteins in live cells, but generating specific intrabodies against the thousands of proteins in a proteome poses a challenge. We leverage the widespread availability of fluorescently labelled proteins to visualize and manipulate intracellular signalling pathways in live cells by using nanobodies targeting fluorescent protein tags.

**Results:** We generated a toolkit of plasmids encoding nanobodies against red and green fluorescent proteins (RFP and GFP variants), fused to functional modules. These include fluorescent sensors for visualization of Ca^2+^, H^+^ and ATP/ADP dynamics; oligomerizing or heterodimerizing modules that allow recruitment or sequestration of proteins and identification of membrane contact sites between organelles; SNAP tags that allow labelling with fluorescent dyes and targeted chromophore-assisted light inactivation; and nanobodies targeted to lumenal sub-compartments of the secretory pathway. We also developed two methods for crosslinking tagged proteins: a dimeric nanobody, and RFP-targeting and GFP-targeting nanobodies fused to complementary hetero-dimerizing domains. We show various applications of the toolkit and demonstrate, for example, that IP_3_ receptors deliver Ca^2+^ to the outer membrane of only a subset of mitochondria, and that only one or two sites on a mitochondrion form membrane contacts with the plasma membrane.

**Conclusions:** This toolkit greatly expands the utility of intrabodies, and will enable a range of approaches for studying and manipulating cell signalling in live cells.

## Background

Visualizing the location of specific proteins within cells and manipulating their function is crucial for understanding cell biology. Antibodies can define protein locations in fixed and permeabilized cells, but antibodies are large protein complexes that are difficult to introduce into live cells [1]. This limits their ability to interrogate the dynamics or affect the function of proteins in live cells. Small protein-based binders, including nanobodies derived from the variable region of the heavy chains (V_HH_) of camelid antibodies, offer a promising alternative [2]. Nanobodies can be encoded by plasmids and expressed in live cells. However, generating nanobodies against thousands of protein variants is daunting, and even for single targets it can be time-consuming, costly and not always successful. A solution to this bottleneck is provided by fluorescently tagged proteins, which are widely used in cell biology [3, 4] after heterologous expression of proteins or gene editing of endogenous proteins [5-7]. The most common application of fluorescent protein (FP) tags is to visualize protein locations, but they have additional potential as generic affinity tags to manipulate and visualize protein functions in live cells. These opportunities are under-developed.

Green fluorescent protein (GFP) has undergone numerous cycles of optimization as a reporter and non-perturbing tag [3, 8]. Most GFP-tagged proteins therefore retain their endogenous localization and function [9]. Large libraries of plasmids encoding GFP-tagged proteins are now available [10]. Proteome-scale expression of GFP-tagged proteins or genome-scale tagging of gene products with GFP has been reported for *Drosophila* [11], fungi [12-14], plants [15, 16] and bacteria [17].

Proteins tagged with red fluorescent proteins (RFPs) such as DsRed, mRFP and mCherry (mCh) are also popular. Extensive optimization has made them attractive tags [3, 18], and libraries of RFP-tagged proteins have been developed in mouse stem cells [19] and yeast [14].

Nanobodies that bind to RFP [20, 21] or GFP [21, 22] are most commonly used in their purified forms for immunoprecipitation and immunocytochemistry. However, they also offer a generic means of targeting in live cells the huge variety of available tagged proteins and the many emerging examples of endogenous proteins tagged with FPs by gene editing. GFP-targeting nanobodies have been used for applications such as targeted proteosomal degradation [23, 24] and relocation of proteins in cells [25], but these and other applications are less developed for RFP-targeting nanobodies.

Here we develop a plasmid-encoded toolkit of nanobodies that bind common FP tags, including RFPs, CFP, GFP and YFP, fused to functional modules for visualization and manipulation of cell signalling (Fig. 1). We fused the nanobodies to a variety of functional modules: fluorescent sensors for Ca^2+^, H^+^ and ATP/ADP; optimized SNAP tags for labelling with bright and photostable dyes [26]; and hetero-dimerizing partners that allow inducible recruitment or sequestration of proteins and visualization of membrane contact sites (MCS) between organelles. We developed two methods to allow crosslinking of RFP-tagged and GFP-tagged proteins: a dimeric nanobody, and co-expression of RFP-targeting and GFP-targeting nanobodies fused to complementary hetero-dimerizing domains. We also describe functionalized nanobodies directed to lumenal sub-compartments of the secretory pathway. We demonstrate the utility of nanobody fusions by visualizing local Ca^2+^ dynamics at the surface of mitochondria, by manipulating the locations of proteins and organelles within cells, by characterizing MCS between mitochondria and the plasma membrane (PM), and by targeting lumenal Ca^2+^ sensors to a sub-compartment of the endoplasmic reticulum (ER).

**Fig 1.**
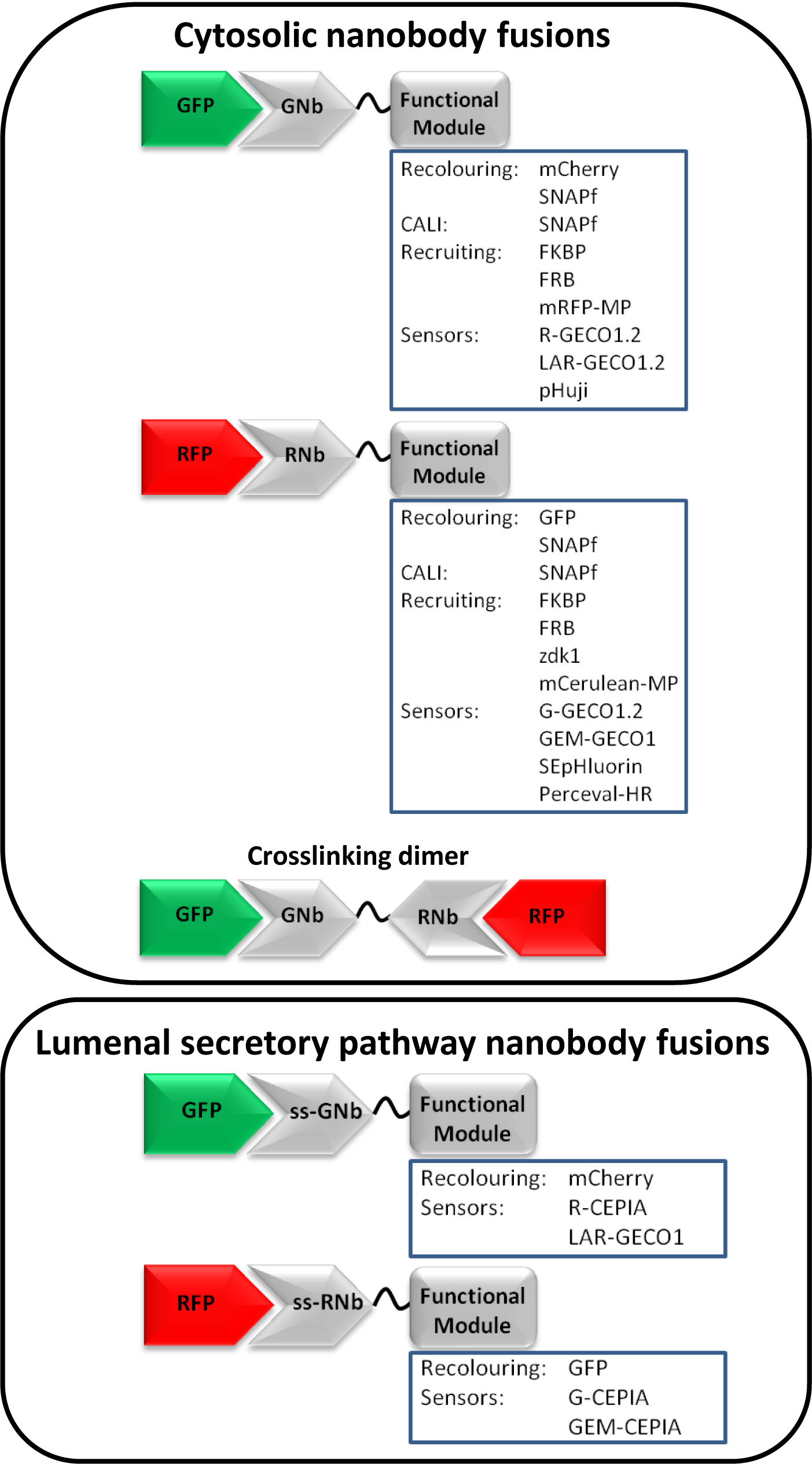
Nanobody fusions for visualizing and manipulating intracellular signalling. Plasmids were generated that encode nanobodies specific for GFP variants (GNb) or RFP variants (RNb), fused to functional modules. Nanobody fusions with an N-terminal signal sequence to target them to the secretory pathway are also shown (ss-GNb and ss-RNb).

This versatile toolkit of genetically-encoded, functionalized nanobodies greatly expands the utility of RFP- and GFP-targeting nanobodies. It will provide a valuable resource for studying protein function and cell signalling in live cells. We illustrate some applications and demonstrate, for example, that IP_3_ receptors deliver Ca^2+^ to the outer membrane of only some mitochondria and that MCS between mitochondria and the plasma membrane occur at only one or two sites on each mitochondrion.

## Results

### Targeting RFP and GFP variants with genetically-encoded nanobody fusions in live cells

The RFP nanobody (RNb) and GFP nanobody (GNb) used are the previously described llama variants LaM4 and LaG16, respectively [21]. They were chosen for their favourable combinations of high affinity (K_d_ values of 0.18 nM and 0.69 nM, respectively) and ability to bind a variety of RFP or GFP variants [21]. The latter attribute maximizes their potential for targeting a wide variety of FPs [3, 4]. LaM4 binds both mCh and DsRed variants, but not GFPs [21]. In addition to binding GFP, LaG16 binds cyan, blue and yellow FPs (CFP, BFP and YFP), but not RFPs [21]. In contrast, the widely used VhhGFP4 nanobody binds GFP, but not CFP [22].

In HeLa cells with organelles (ER, mitochondria, nucleus and lysosomes) labelled with mCh or mRFP markers, expression of RNb-GFP (Fig. 2A) specifically identified the labelled organelle (Fig. 2B). Similar results were obtained with GNb-mCh (Fig. 2C) and organelles (ER, mitochondria and nucleus) labelled with GFP or mTurquoise (Fig. 2D). These results demonstrate that plasmid-encoded RNb and GNb allow specific labelling of a variety of RFP and GFP variants in live cells.

**Fig 2.**
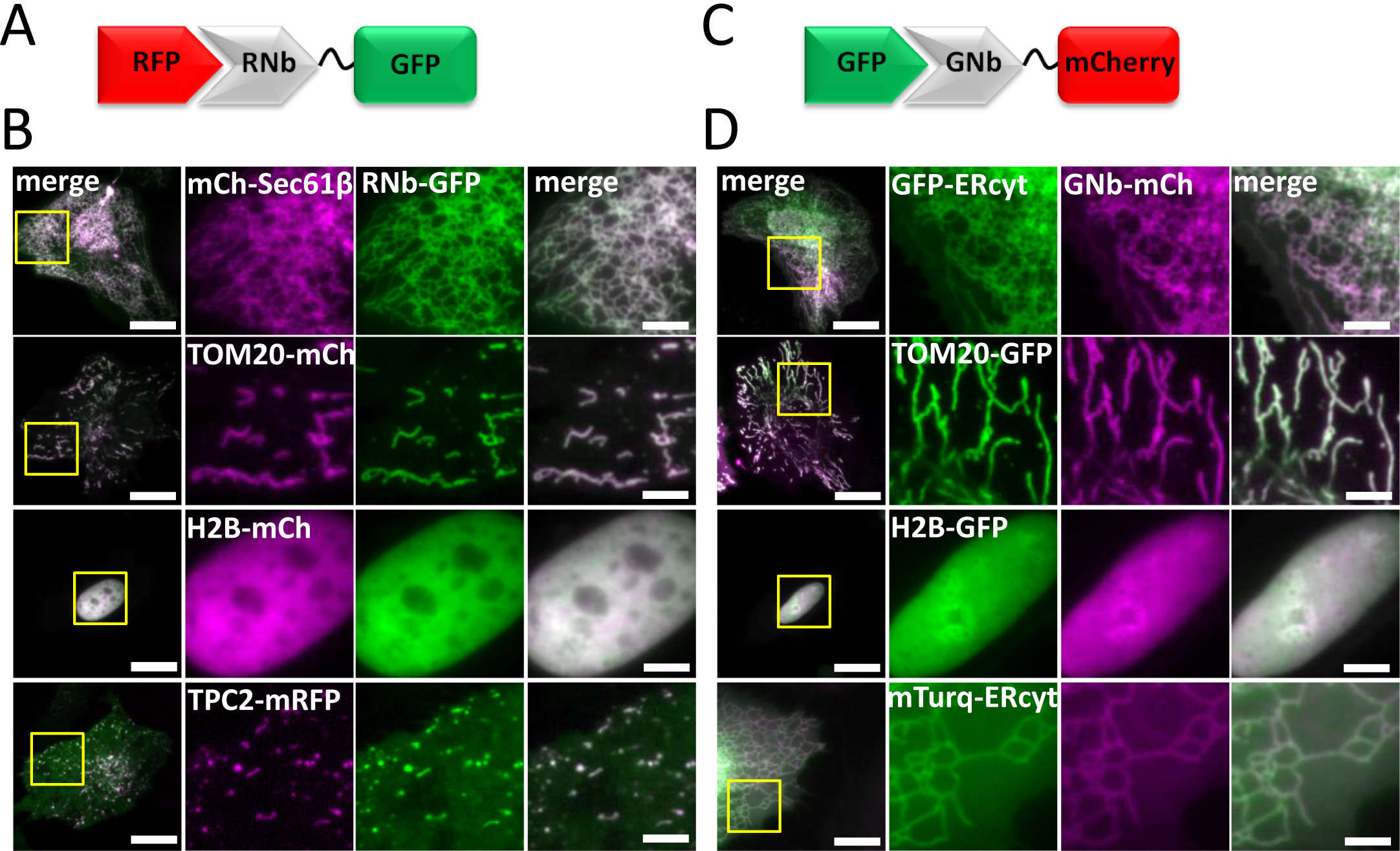
RNb and GNb fusion proteins bind to their respective tagged proteins in live cells. (**A**) Schematic of the RNb-GFP fusion binding to RFP. (**B**) HeLa cells expressing RNb-GFP with RFP-tagged markers for the ER surface (mCh-Sec61β), the mitochondrial surface (TOM20-mCh), the nucleus (H2B-mCh) or the surface of lysosomes (TPC2-mRFP). Cells were imaged in HBS using epifluorescence microscopy (cells expressing H2B-mCh) or TIRFM (other cells). Yellow boxes indicate regions enlarged in the subsequent panels. Colocalization values (Pearson’s coefficient, *r*) were: mCh-Sec61β (*r* = 0.93 ± 0.09, n = 10 cells); TOM20-mCh (*r* = 0.94 ± 0.09, n = 10 cells); H2B-mCh (*r* = 0.97 ± 0.06, n = 10 cells) and TPC2-mRFP (*r* = 0.78 ± 0.09, n = 5 cells). (**C**) Schematic of the GNb-mCh fusion binding to GFP. (**D**) HeLa cells co-expressing GNb-mCh with GFP-tagged markers for the ER surface (GFP-ERcyt), the mitochondrial surface (TOM20-GFP) and the nucleus (H2B-GFP), or an mTurquoise2-tagged ER-surface marker (mTurq-ERcyt). Cells were imaged using epifluorescence microscopy (cells expressing H2B-GFP) or TIRFM (other cells). Yellow boxes indicate regions enlarged in the subsequent panels. Colocalization values were: GFP-ERcyt (*r* = 0.92 ± 0.08, n = 8 cells); TOM20-GFP (*r* = 0.87 ± 0.05, n = 7 cells); H2B-GFP (*r* = 0.94 ± 0.07, n = 6 cells) and mTurq-ERcyt (*r* = 0.97 ± 0.03, n = 7 cells). Scale bars 10 µm (main images) or 2.5 µm (enlargements).

### Targeting sensors to RFP and GFP

The effects of intracellular messengers such as Ca^2+^ [27], H^+^ [28] and ATP/ADP [29] can be highly localized within cells. To enable visualization of these intracellular messengers in microdomains around RFP-tagged and GFP-tagged proteins, we fused RNb and GNb to fluorescent sensors for Ca^2+^ [30], H^+^ [31, 32] or ATP/ADP [33].

RNb was fused to the green fluorescent Ca^2+^ sensor G-GECO1.2 (Fig. 3), and GNb was fused to the red fluorescent Ca^2+^ sensors, R-GECO1.2 or LAR-GECO1.2 [30] (Fig. 4). The affinities of these sensors for Ca^2+^ (of 1.2 µM for G-GECO1.2 and R-GECO1.2, and 10 µM for LAR-GECO1.2) are low relative to global changes in the cytosolic free Ca^2+^ concentration ([Ca^2+^]_c_) after receptor stimulation (typically ~300 nM) [34]. This facilitates selective detection of the large, local rises in [Ca^2+^] that are important for intracellular signalling, at the contacts between active inositol 1,4,5-trisphosphate receptors (IP_3_Rs) and mitochondria, for example [27]. To allow targeted measurement of relatively low resting [Ca^2+^] within cellular microdomains we also fused RNb to the ratiometric Ca^2+^-sensor, GEM-GECO1 = 300 nM) [30], to give RNb-GEMGECO1 (***Additional file 1: Fig. S1***).

**Fig 3.**
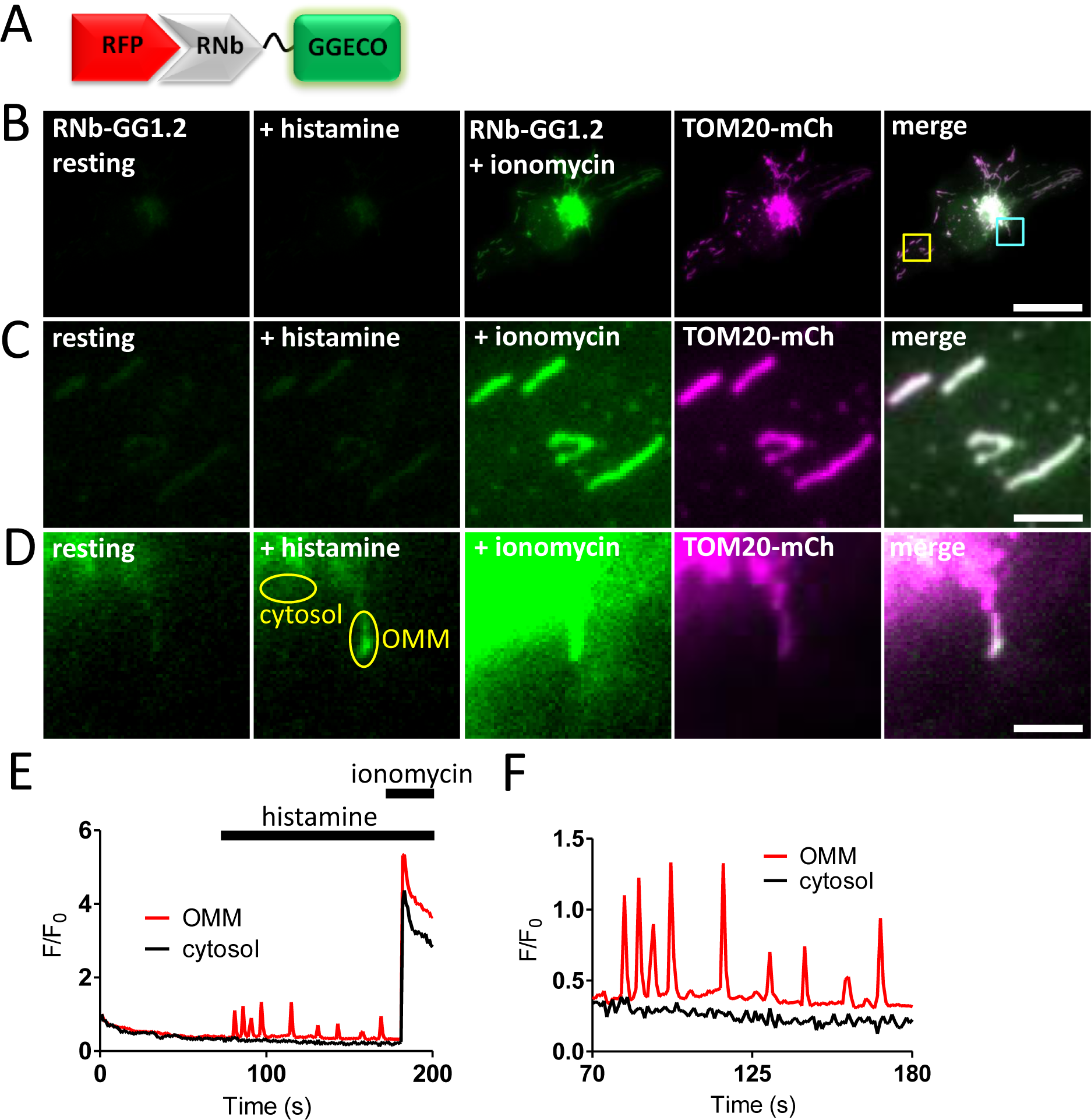
Targeting RNb-Ca^2+^ sensors to RFP-tagged proteins. (**A**) Schematic of RNb-GGECO fusion binding to RFP. (**B-D**) HeLa cells expressing RNb-GGECO1.2 and TOM20-mCh, before and after addition of histamine (100 µM) and then ionomycin (5 µM). Cells were imaged in HBS using TIRFM. The TOM20-mCh image is shown after the histamine and ionomycin additions. The merged images are shown using images of RNb-GGECO1.2 after ionomycin (B, C) or histamine (D). The yellow and cyan boxed regions in panel B are shown enlarged in panels C and D, respectively. Scale bars are 10 µm (B) or 1.25 µm (C, D). (**E**) Timecourse of the effects of histamine (100 μM) and ionomycin (5 μM) on the fluorescence of RNb-GGECO1.2 (F/F_0_, where F and F_0_ are fluorescence recorded at t and t = 0). The traces are from regions coinciding with a single mitochondrion or cytosol (regions identified in panel D), indicating changes in [Ca^2+^] at the OMM. (**F**) Enlarged region (70-180 s) of the graph shown in E. Results are representative of cells from 13 independent experiments.

**Fig 4.**
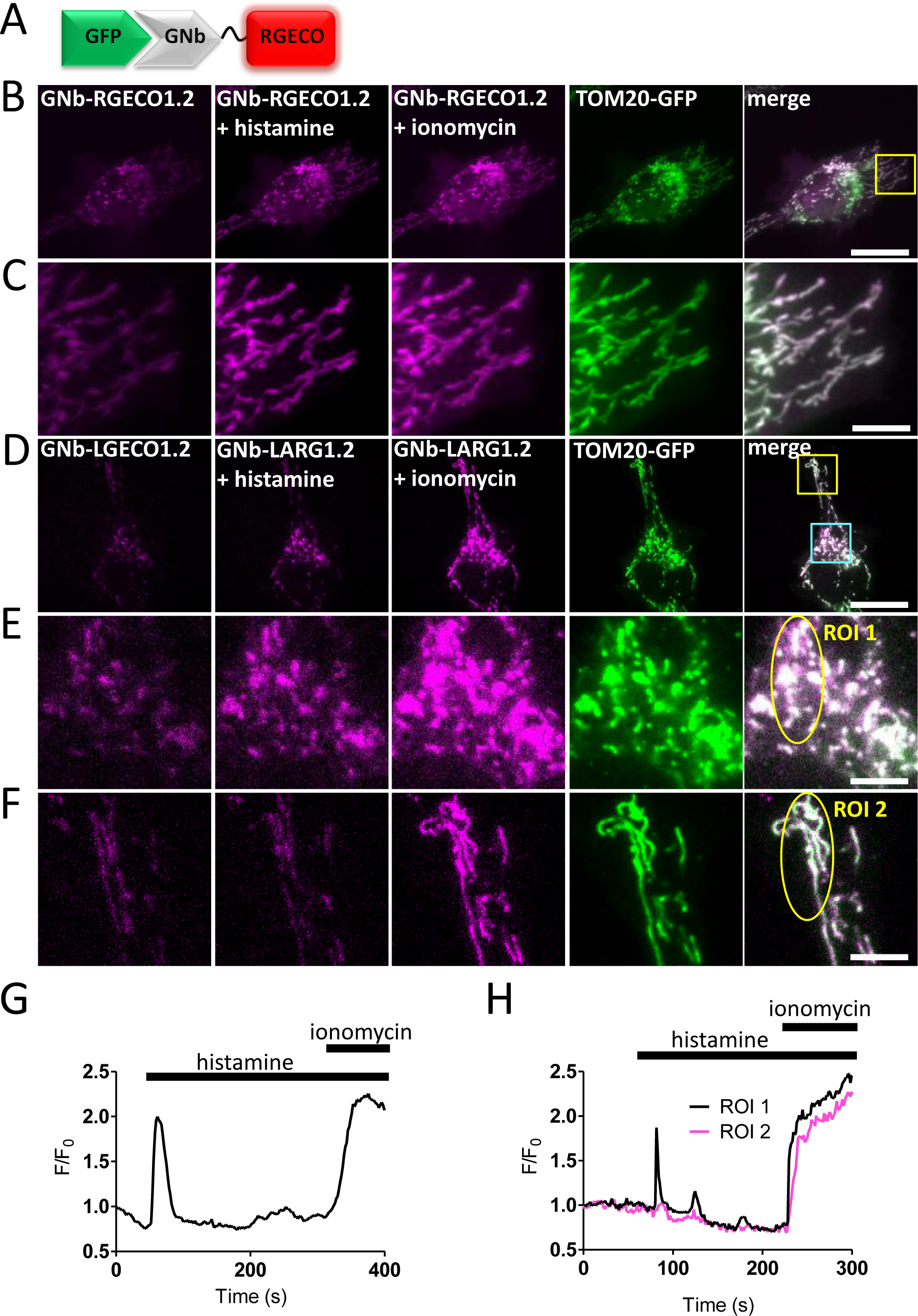
Targeted GNb-Ca^2+^ sensors detect changes in [Ca^2+^] at the surface of mitochondria. (**A**) Schematic of GNb-RGECO fusion binding to GFP. (**B, C**) Representative HeLa cells co-expressing TOM20-GFP and GNb-RGECO1.2 imaged in HBS using TIRFM before and after addition of histamine (100 µM) and then ionomycin (5 µM). The TOM20-GFP images are shown after the histamine and ionomycin additions. Histamine and ionomycin evoked changes in fluorescence of GNb-RGECO1.2 at the OMM. The yellow boxed region in panel B is shown enlarged in panel C. (**D-F**) Similar analyses of HeLa cells co-expressing TOM20-GFP and GNb-LAR-GECO1.2 (GNb-LARG1.2). Histamine (100 μM) evoked changes in fluorescence of GNb-LARG1.2 at the OMM of mitochondria in the perinuclear region (region of interest 1 (ROI 1) in E), but not in a peripheral region (ROI 2 in F). All mitochondria responded to ionomycin (5 μM), indicating that histamine evoked local changes in [Ca^2+^] at the OMM. The cyan and yellow boxed regions in D are shown enlarged in E and F, respectively. Scale bars 10 μm (B, D) or 2.5 μm (C, E, F). (**G**) Timecourse of the changes in fluorescence of GNb-RGECO1.2 at the OMM evoked by histamine and ionomycin for the entire cell shown in B. (**H**) Fluorescence changes recorded from ROI 1 and ROI 2 in panels E and F. Results are representative of cells from 4 independent experiments.

In HeLa cells expressing TOM20-mCh or TOM20-GFP to identify the outer mitochondrial membrane (OMM), the RNb-Ca^2+^ sensors (Fig. 3 and ***Additional file 1: Fig. S1***) and GNb-Ca^2+^ sensors (Fig. 4) were targeted to the OMM. Both families of targeted sensor reported an increase in [Ca^2+^] after treatment with the Ca^2+^ ionophore, ionomycin (Fig. 3, Fig. 4 and ***Additional file 1: Fig. S1***). This confirms the ability of the sensors to report [Ca^2+^] changes when targeted to the OMM microdomain.

In some cells, the targeted Nb-Ca^2+^ sensors revealed local changes in [Ca^2+^]_c_ after receptor stimulation with histamine, which stimulates IP_3_ formation and Ca^2+^ release from the ER in HeLa cells [34]. Imperfect targeting of the RNb-GGECO1.2 to the OMM allowed Ca^2+^ signals at the surface of individual mitochondria to be distinguished from those in nearby cytosol in some cells (Fig. 3D-F and ***Additional file 2: Video 1***). In the example shown, RNb-GGECO1.2 at both the OMM and nearby cytosol responded to the large, global increases in [Ca^2+^] evoked by ionomycin. However, cytosolic RNb-GGECO1.2 did not respond to histamine, while the sensor at the OMM responded with repetitive spiking (Fig. 3D-F and ***Additional file 2: Video 1***). The GNb-LARGECO1.2 sensor, which has the lowest affinity for Ca^2+^ of the sensors used, revealed changes in [Ca^2+^]_c_ at the surface of some mitochondria, but not others in the same cell (Fig. 4D-F, Fig. 4H and ***Additional file 3: Video 2***). In the example shown, GNb-LARGECO1.2 at the OMM in all mitochondria within the cell responded to the large, global increases in [Ca^2+^] evoked by ionomycin. However, in response to histamine mitochondria in the perinuclear region responded, but not those in peripheral regions (Fig. 4D-F, Fig. 4H and ***Additional file 3: Video 2***). Ca^2+^ uptake by mitochondria affects many cellular responses, including mitochondrial metabolism, ATP production and apoptosis [35]; and Ca^2+^ at the cytosolic face of the OMM regulates mitochondrial motility [36]. The subcellular heterogeneity of mitochondrial exposure to increased [Ca^2+^] suggests that these responses may be very localized in cells.

These observations align with previous reports showing that Ca^2+^-mobilizing receptors evoke both oscillatory [Ca^2+^] changes within the mitochondrial matrix [37], and large local increases in [Ca^2+^] at the cytosolic face of the OMM [38]. Our results establish that nanobody-Ca^2+^-sensor fusions are functional and appropriately targeted, and can be used to detect physiological changes in [Ca^2+^] within cellular microdomains such as the OMM.

For targeted measurements of intracellular pH, RNb was fused to the green fluorescent pH sensor super-ecliptic pHluorin (SEpHluorin) [31], and GNb was fused to the red fluorescent pH sensor pHuji [32]. Both Nb-pH sensors were targeted to the OMM by the appropriate fluorescent tags, where they responded to changes in intracellular pH imposed by altering extracellular pH in the presence of the H^+^/K^+^ ionophore nigericin (Fig. 5).

**Fig 5.**
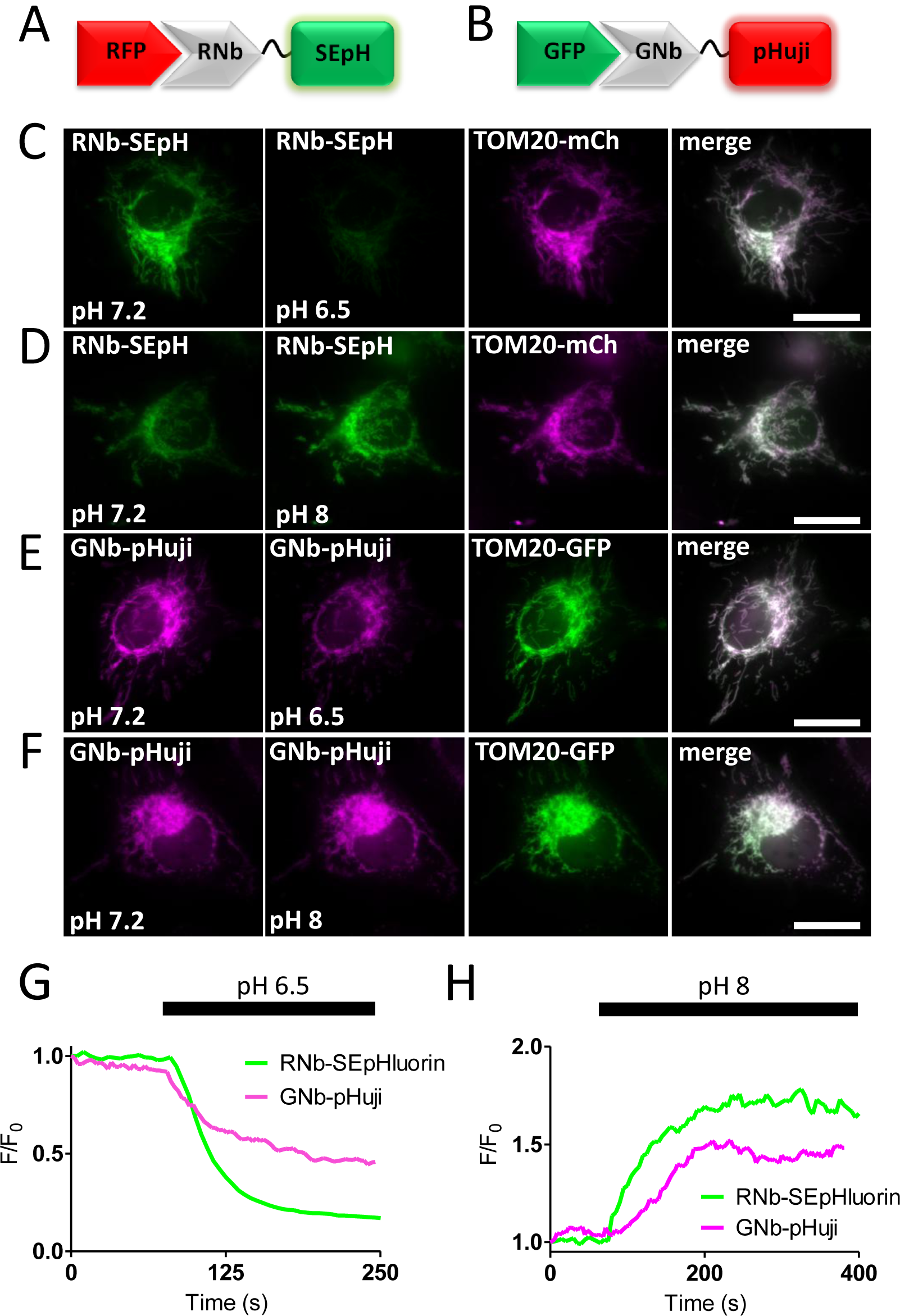
Targeting H^+^ sensors to RFP-tagged and GFP-tagged proteins. (**A**) Schematic of RNb fused to the pH sensor superecliptic pHluorin (RNb-SEpH) and bound to RFP. (**B**) Schematic of GNb-pHuji binding to RFP. (**C**, **D**) HeLa cells co-expressing RNb-SEpH and TOM20-mCh were imaged in modified HBS (MHBS) using epifluorescence microscopy and exposed to extracellular pH 6.5 (C) or pH 8 (D) in the presence of nigericin (10 µM). Scale bars 10 μm. (**E, F**) HeLa cells co-expressing GNb-pHuji and TOM20-GFP were exposed to extracellular pH 6.5 (E) or pH 8 (F) in the presence of nigericin. Scale bars 10 μm. (**G**, **H**) Timecourse from single cells of the fluorescence changes (F/F_0_) of mitochondrially targeted RNb-SEpH or GNb-pHuji evoked by the indicated manipulations of extracellular pH. Results shown are representative of 3 independent experiments.

For targeted measurements of ATP/ADP, RNb was fused to the excitation-ratiometric ATP/ADP sensor Perceval-HR [33]. RNb-Perceval-HR was targeted to the surface of mitochondria and responded to inhibition of glycolysis and oxidative phosphorylation (Fig. 6).

**Fig 6.**
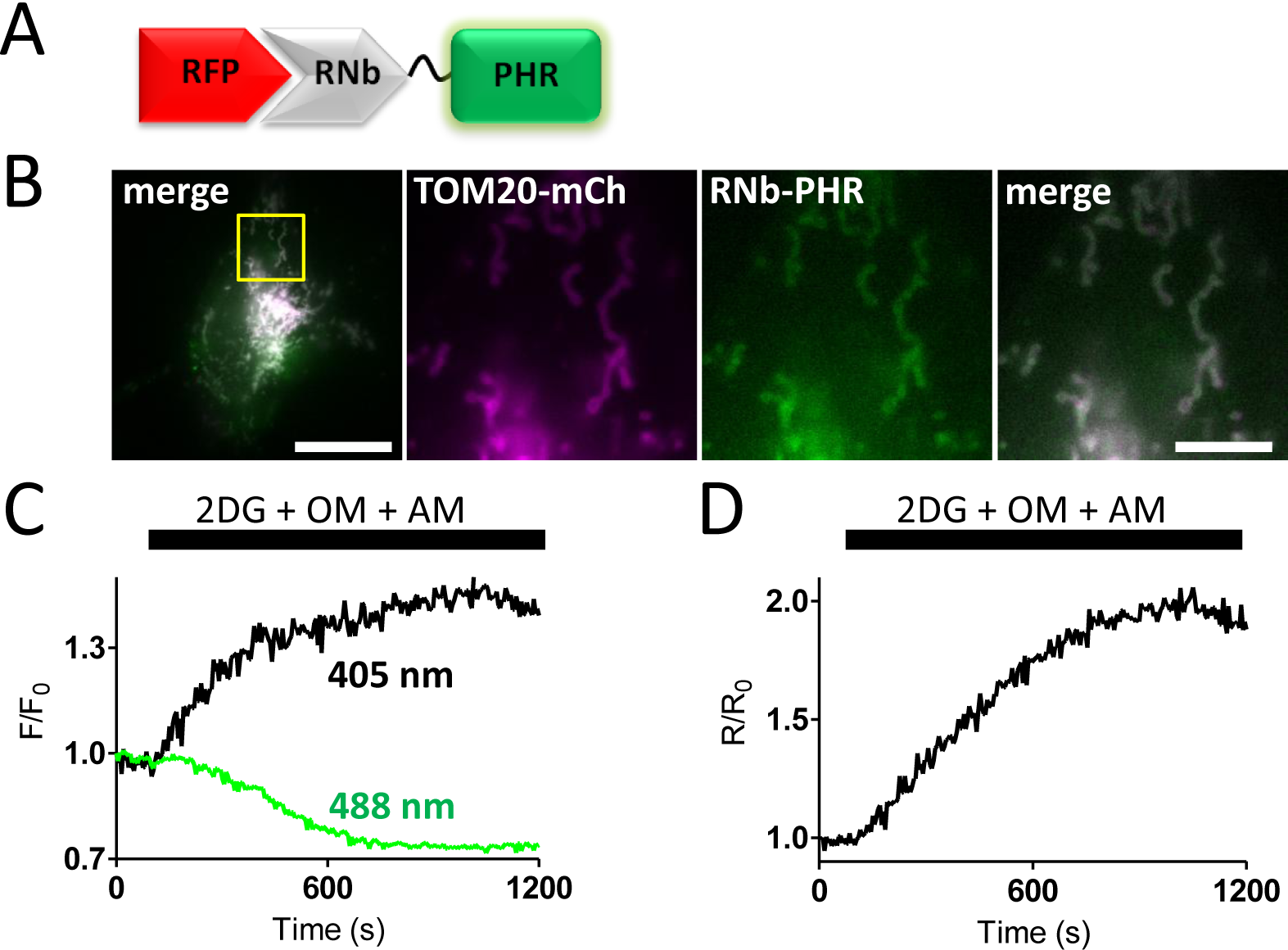
Targeting an ATP/ADP sensor to RFP-tagged proteins. (**A**) Schematic of RNb-Perceval-HR fusion (RNb-PHR) bound to RFP. (**B**) HeLa cells co-expressing RNb-PHR and TOM20-mCh were imaged in HBS using epifluorescence microscopy. The yellow box indicates the region enlarged in subsequent panels. Scale bars 10 μm (main image) and 2.5 μm (enlarged images). (**C, D**) Changes in fluorescence for each excitation wavelength (405 and 488 nm, F/F_0_) (C) and their ratio (R/R_0_, where R = F_405_/F_488_) (D) of mitochondrially targeted RNb-Perceval-HR after addition of 2-deoxyglucose (2DG, 10 mM), oligomycin (OM, 1 µM) and antimycin (AM, 1 μM). The results indicate a decrease in the ATP/ADP ratio at the OMM. Results are representative of 3 independent experiments.

The results demonstrate that nanobodies can be used to direct sensors for Ca^2+^, H^+^ or ATP/ADP to specific subcellular compartments tagged with variants of RFP or GFP.

### Targeting SNAPf tags to RFP and GFP in live cells

SNAP, and related tags, are versatile because a range of SNAP substrates, including some that are membrane-permeant, can be used to attach different fluorophores or cargoes to the tag [39]. Purified GFP-targeting nanobodies fused to a SNAP tag have been used to label fixed cells for optically demanding applications [40]. We extended this strategy to live cells using RNb and GNb fused to the optimized SNAPf tag [41] (Fig. 7A and B). In cells expressing the mitochondrial marker TOM20-mCh, RNb-SNAPf enabled labelling of mitochondria with the cell-permeable substrate SNAP-Cell 647-SiR and imaging at far-red wavelengths (Fig. 7C). In cells expressing lysosomal LAMP1-mCh and RNb-SNAPf, SNAP-Cell 647-SiR instead labelled lysosomes (Fig. 7D), demonstrating that SNAP-Cell 647-SiR specifically labelled the organelles targeted by RNb-SNAPf. Similar targeting of SNAP-Cell 647-SiR to mitochondria (Fig. 7E) and lysosomes (Fig. 7F) was achieved by GNb-SNAPf co-expressed with the appropriate GFP-tagged organelle markers.

**Fig 7.**
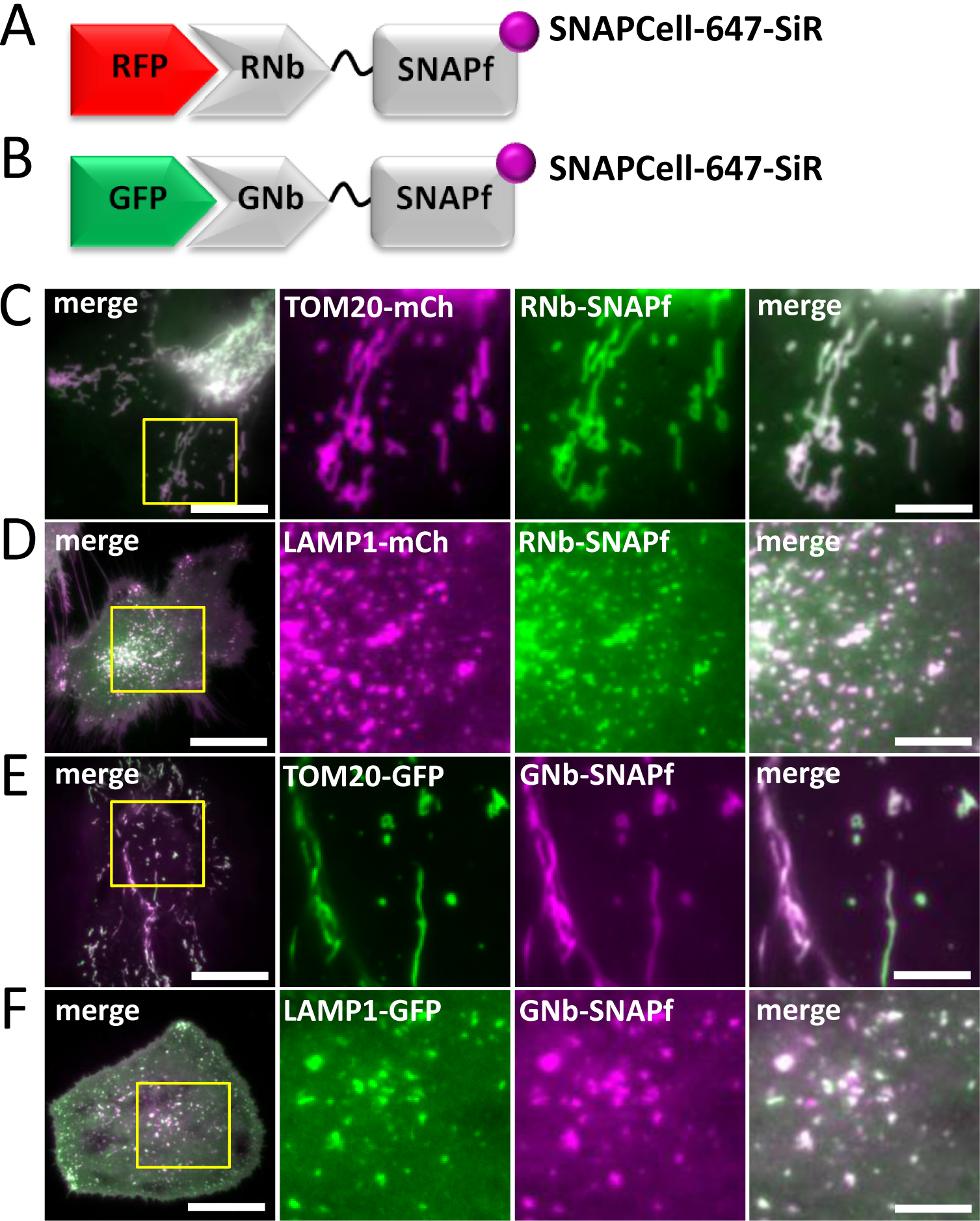
Nanobody-SNAPf fusion proteins allow labelling of RFP-tagged and GFP-tagged proteins with fluorescent O^6^-benzylguanine derivatives in live cells. (**A, B**) Schematics of RNb-SNAPf fusion bound to RFP, and GNb-SNAPf fusion bound to GFP, after labelling with SNAP-Cell-647-SiR (magenta circles). (**C-F**) HeLa cells co-expressing RNb-SNAPf and mitochondrial TOM20-mCh (C), RNb-SNAPf and lysosomal LAMP1-mCh (D), GNb-SNAPf and TOM20-GFP (E) or GNb-SNAPf and LAMP1-GFP (F) were treated with SNAP-Cell-647-SiR (0.5 µM, 30 min at 37°C) and imaged using TIRFM. Scale bars 10 µm (main images) or 2.5 µm (enlarged images of yellow boxed regions). Colocalization values: RNb-SNAPf + TOM20-mCh (*r* = 0.95 ± 0.02, n = 6 cells); RNb-SNAPf + LAMP1-mCh (*r* = 0.84 ± 0.06, n = 8 cells); GNb-SNAPf + TOM20-GFP (*r* = 0.78 ± 0.09, n = 10 cells); and GNb-SNAPf + LAMP1-GFP (*r* = 0.85 ± 0.10, n = 11 cells).

Chromophore-assisted light inactivation (CALI) can inactivate proteins or organelles by exciting fluorophores attached to them that locally generate damaging reactive superoxide. Historically, antibodies were used to direct a photosensitizer to its target, but fusion of fluorescent proteins or SNAP tags to proteins of interest is now widely used [42]. RNb-SNAPf and GNb-SNAPf make the SNAP strategy more broadly applicable to CALI applications. We demonstrate this by targeting CALI to the outer surface of lysosomes. We anticipated that CALI in this microdomain might, amongst other effects, disrupt the motility of lysosomes, which depends on their association with molecular motors [43]. RNb-SNAPf enabled labelling of lysosomes with the CALI probe fluorescein, using the cell-permeable substrate, SNAP-Cell-fluorescein (Fig. 8A and B). Exposure to blue light then immobilized the lysosomes (Fig. 8C-F and ***Additional file 4: Video 3***), indicating a loss of motor-driven motility. Control experiments demonstrated that labelling cytosolic SNAPf with SNAP-Cell-fluorescein (***Additional file 1: Fig. S2A*** and ***S2B***) had significantly less effect on lysosomal motility after exposure to blue light (Fig. 8F and ***Additional file 1: Fig. S2C-E***). These results demonstrate that nanobody-SNAPf fusions allow targeting of fluorescent dyes in live cells, which can be used for re-colouring of tagged proteins or targeted CALI.

**Fig 8.**
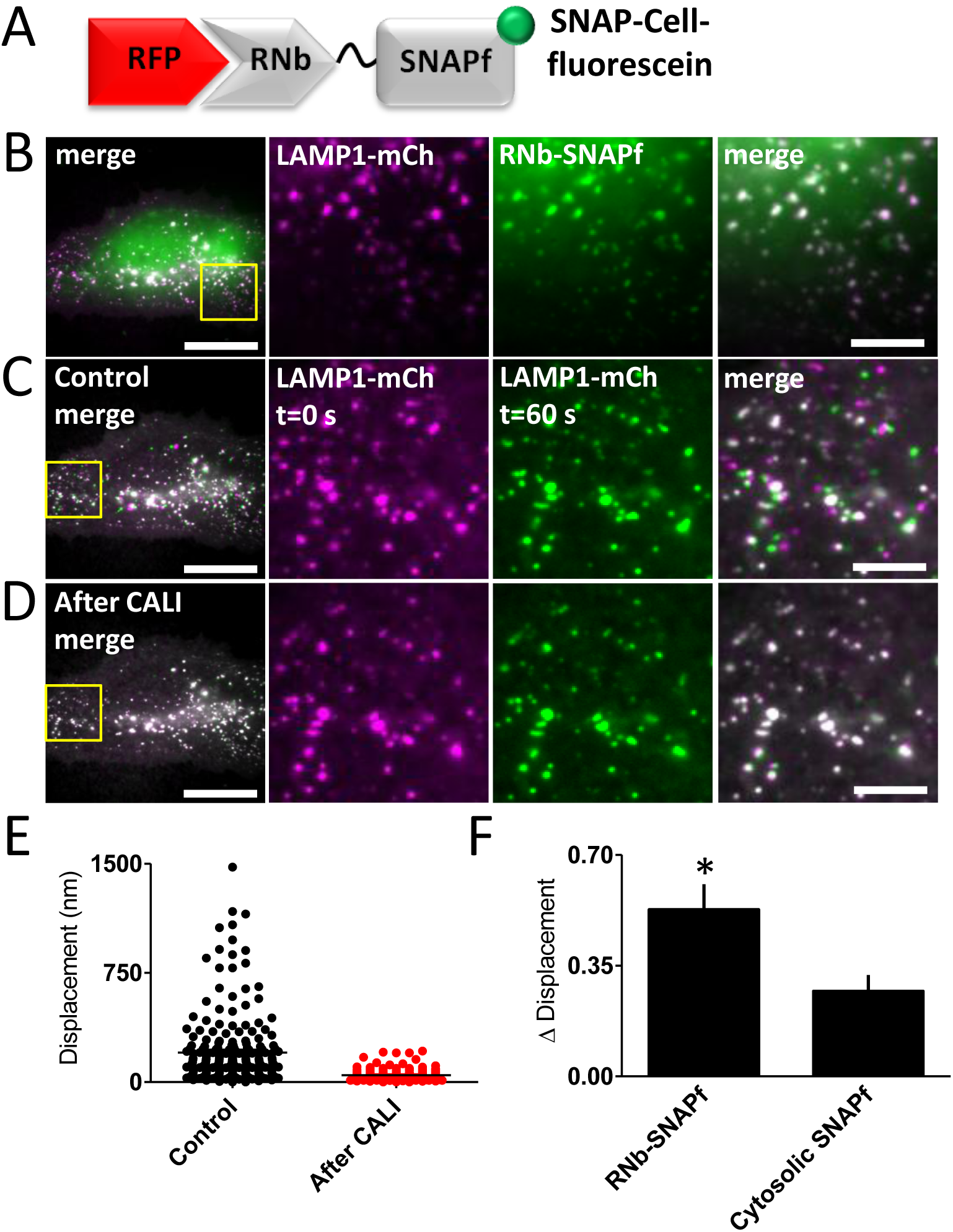
Targeting CALI to lysosomes using RNb-SNAPf reduces lysosomal motility. (**A**) Schematic of RNb-SNAPf after labelling with SNAP-Cell-fluorescein (green circle) and bound to RFP. (**B**) HeLa cells co-expressing LAMP1-mCh and RNb-SNAPf were incubated with SNAP-Cell-fluorescein (0.5 µM, 30 min, 37°C), which labelled lysosomes (colocalization values, *r* = 0.73 ± 0.02, n = 6 cells), and imaged using TIRFM. (**C, D**) Cells were then exposed to 488-nm light for 3 s to induce CALI. TIRFM images show a representative cell at different times before (C) and after (D) CALI, with the image at t = 0 s shown in magenta and the image at t = 60 s in green. White in the merged images from the two different times indicates immobile lysosomes, while green and magenta indicate lysosomes that moved in the interval between images. Yellow boxes show regions enlarged in subsequent images. Scale bars 10 µm (main images) and 2.5 μm (enlargements). For clarity, images were auto-adjusted for brightness and contrast (ImageJ) to compensate for bleaching of mCh during tracking and CALI. (**E**) Effect of CALI on the displacements of individual lysosomes, determining by particle-tracking (TrackMate), during a 60-s recording from a representative cell (images taken every 1 s; mean values shown by bars). (**F**) Summary data (mean ± SEM, n = 6 cells from 6 independent experiments) show the mean fractional decrease in displacement (Δ Displacement) due to CALI in cells expressing RNb-SNAPf or cytosolic SNAPf (see ***Additional file 1: Fig. S2***). The fractional decrease in displacement for each cell was defined as: (MD_pre_ – MD_post_) / MD_pre_, where MD_pre_ and MD_post_ are the mean displacement of all tracked particles in 60 s before and after CALI. \**p* < 0.05, unpaired Student’s *t*-test.

### Sequestration of proteins tagged with RFP or GFP

Fusion of GFP nanobodies to degrons allows proteosomal degradation of GFP-tagged proteins [24], but the method is slow and cumbersome to reverse. An alternative strategy is to sequester tagged proteins so they cannot fulfil their normal functions. We used two strategies to achieve this: artificial clustering and recruitment to mitochondria.

We induced artificial clustering by fusing RNb or GNb to a multimerizing protein (MP) comprising a dodecameric fragment of Ca^2+^-calmodulin-dependent protein kinase II (CaMKII) [44], with an intervening fluorescent protein (mRFP or mCerulean) for visualization of the Nb fusion (Fig. 9A and B). RNb-mCerulean-MP caused clustering of the ER transmembrane protein mCh-Sec61β (Fig. 9C and D) and caused lysosomes tagged with LAMP1-mCh to aggregate into abnormally large structures (Fig. 9E and F). GNb-mRFP-MP had the same clustering effect on lysosomes labelled with LAMP1-GFP (Fig. 9G and H) and caused clustering of GFP-tagged proteins in the cytosol (calmodulin, Fig. 9I and J), nucleus and cytosol (p53, Fig. 9K and L) or ER membranes (IP_3_R3, Fig. 9M and N).

**Fig 9.**
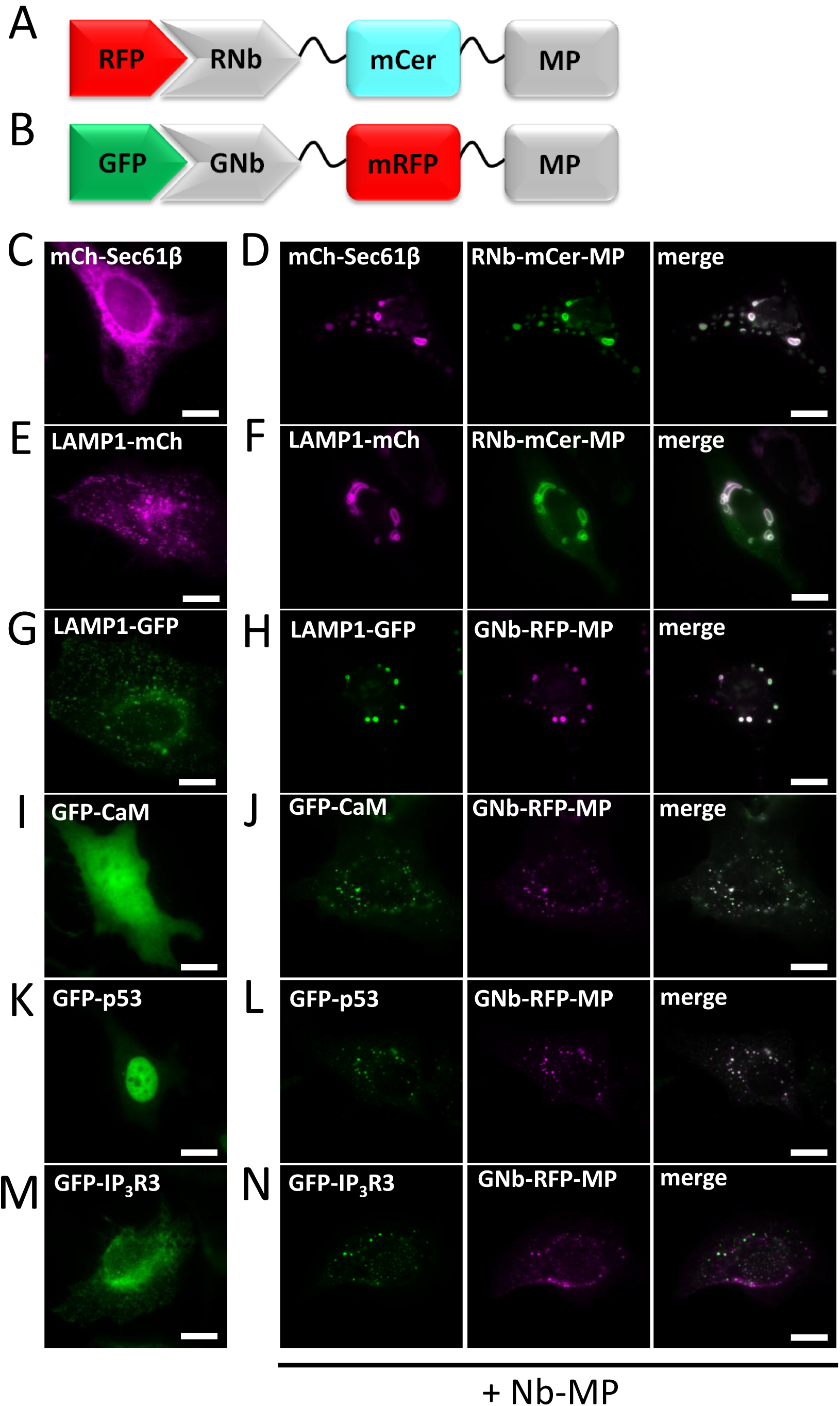
Clustering of RFP-tagged and GFP-tagged proteins and organelles using RNb-mCerulean-MP and GNb-mRFP-MP. (**A**) Schematic of RNb-mCerulean-MP fusion bound to RFP. (**B**) Schematic of GNb-mRFP-MP fusion bound to GFP. (**C-F**) HeLa cells expressing RFP-tagged proteins in the absence (**C, E**) or presence (**D**, **F**) of co-expressed RNb-mCerulean-MP (RNb-mCer-MP) were imaged using epifluorescence microscopy. (**G-N**) HeLa cells expressing GFP-tagged proteins in the absence (G, I, K, M) or presence (H, J, L, N) of co-expressed GNb-mRFP-MP were imaged using epifluorescence microscopy. Results are representative of at least 5 cells, from at least 3 independent experiments. Scale bars 10 µm.

For inducible sequestration, sometimes known as ‘knocksideways’ [45], we used two approaches based on hetero-dimerizing modules, one chemical and one optical. First, we adapted the original knocksideways method, where proteins tagged with FKBP (FK506-binding protein) are recruited by rapamycin to proteins tagged with FRB (FKBP-rapamycin-binding domain) on the OMM, and thereby sequestered. The method has hitherto relied on individual proteins of interest being tagged with FKBP [45]. RNb-FKBP and GNb-FKBP (Fig. 10A and B) extend the method to any protein tagged with RFP or GFP. For our analyses, we expressed TOM70 (an OMM protein) linked to FRB through an intermediary fluorescent protein (GFP or mCh, to allow optical identification of the fusion protein). RNb-FKBP sequestered the ER transmembrane protein mCh-Sec61β at the OMM (TOM70-GFP-FRB) within seconds of adding rapamycin (***Additional file 5: Video 4***) and rapidly depleted mCh-Sec61β from the rest of the ER (Fig. 10C-E). After addition of rapamycin, GNb-FKBP rapidly sequestered endogenous IP_3_R1 tagged with GFP (GFP-IP_3_R1) [7] (Fig. 10F and G, and ***Additional file 6: Video 5***) and cytosolic GFP-tagged calmodulin (Fig. 10H and ***Additional file 7: Video 6***) at mitochondria expressing TOM70-mCh-FRB. Rapamycin caused no sequestration in the absence of the nanobody fusions (***Additional file 1: Fig. S3***).

**Fig 10.**
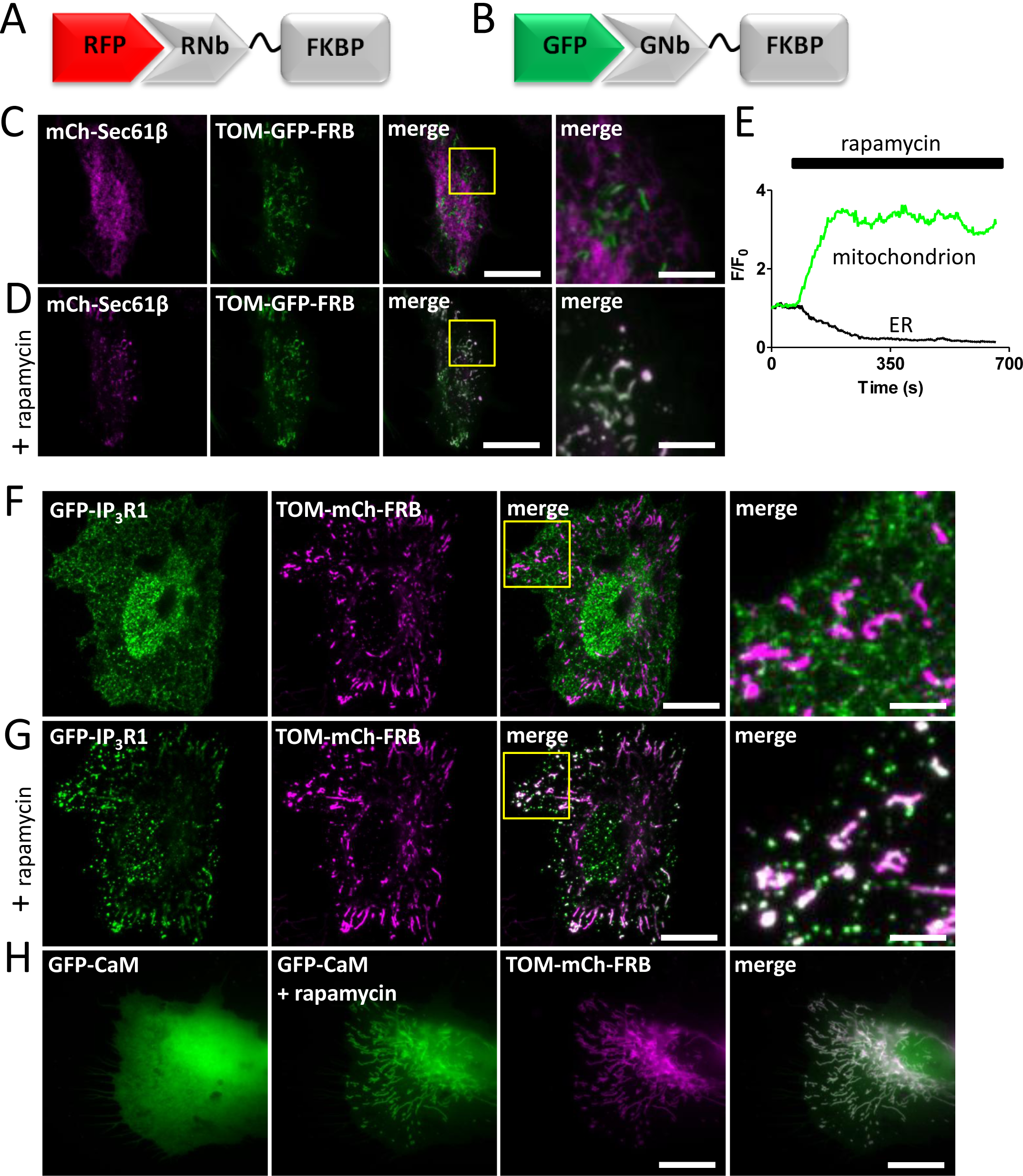
RNb-FKBP inducibly recruits ER transmembrane proteins to mitochondria. (**A**) Schematic of RNb-FKBP bound to RFP. (**B**) Schematic of GNb-FKBP bound to GFP. (**C, D**) HeLa cells co-expressing RNb-FKBP, mitochondrial TOM70-GFP-FRB and mCh-Sec61β were imaged using TIRFM. A representative cell (n = 7) is shown before (C) and after (D) treatment with rapamycin (100 nM, 10 min). The boxed region is enlarged in subsequent images. Scale bars 10 µm (main images) and 2.5 μm (enlargements). (**E**) Timecourse of mCh-Sec61β fluorescence changes (F/F_0_) evoked by rapamycin recorded at a representative mitochondrion and in nearby reticular ER. Results show ~80% loss of fluorescence from the ER devoid of mitochondrial contacts. (**F**, **G**) HeLa cells co-expressing endogenously tagged GFP-IP_3_R1, GNb-FKBP and mitochondrial TOM70-mCh-FRB were imaged using TIRFM. A representative cell (n = 6) is shown before (F) and after (G) treatment with rapamycin (100 nM, 10 min). The boxed region is enlarged in subsequent images. Scale bars 10 µm (main images) and 2.5 μm (enlargements). (**H**) HeLa cells co-expressing GFP-calmodulin (GFP-CaM), GNb-FKBP and TOM20-mCh-FRB were imaged using epifluorescence microscopy. A representative cell (n = 3) is shown before and after treatment with rapamycin (100 nM, 10 min). The image for TOM-mCh-FRB is shown in the presence of rapamycin. Scale bar 10 μm.

To make sequestration reversible and optically activated, we adapted the light-oxygen-voltage-sensing domain (LOV2)/Zdark (zdk1) system in which light induces dissociation of LOV2-zdk1 hetero-dimers [46]. Because this system is operated by blue light at intensities lower than required for imaging GFP [46], it is most suitable for use with red fluorescent tags. RNb-zdk1 (Fig. 11A) sequestered cytosolic mCh on the OMM in cells expressing TOM20-LOV2, and blue laser light rapidly and reversibly redistributed mCh to the cytosol (Fig. 11B and C).

**Fig 11.**
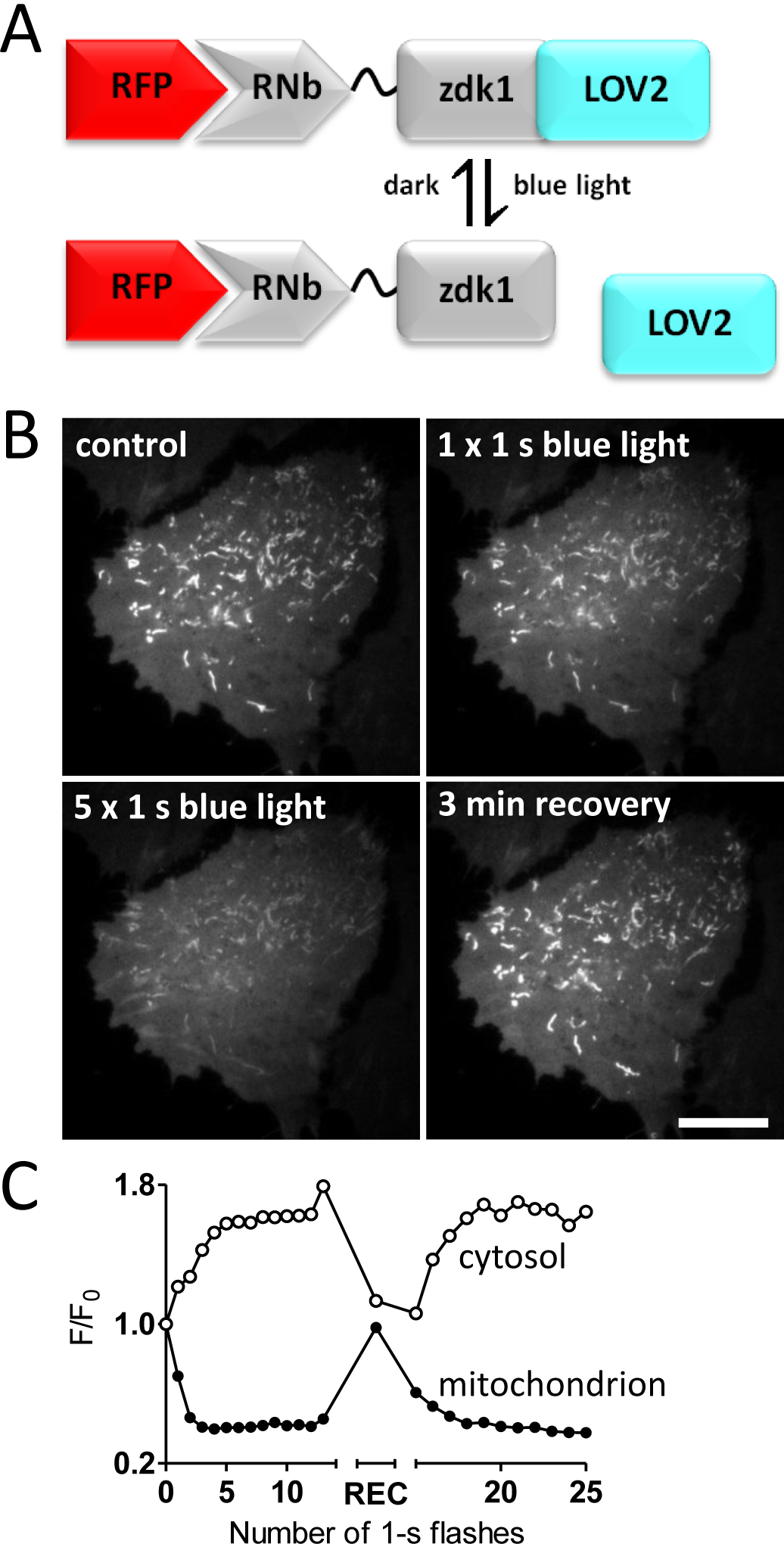
Reversible optogenetic recruitment of RFP-tagged proteins using RNb-zdk1. (**A**) Schematic of RNb-zdk1 fusion bound to RFP, showing the reversible light-evoked dissociation of zdk1 from LOV2. (**B**) HeLa cells co-expressing RNb-zdk1, mitochondrial TOM20-LOV2 and cytosolic mCh were imaged using TIRFM. A representative cell is shown before and after one or five 1-s exposures to blue light (488-nm laser at 2-s intervals) and after a 3-min recovery period in the dark. Scale bar 10 µm. (**C**) Timecourse of the mCherry fluorescence changes (F/F_0_) recorded at a representative mitochondrion and in nearby cytosol after each of the indicated light flashes. There is a reversible decrease (~60%) in mitochondrial mCh fluorescence and a corresponding reversible increase (~70%) in cytosolic fluorescence. A single measurement of mCh fluorescence was made at the end of a 3-min recovery period in the dark (REC) before further light flashes. Results are representative of 5 cells from 3 independent experiments.

### Inducible recruitment of tagged proteins to membrane contact sites

The ability of Nb-FKBP fusions to recruit membrane proteins to FRB-tagged targets suggested an additional application: revealing contact sites between membrane-bound organelles. ER-mitochondrial membrane contact sites (MCS) have been much studied [47], but contacts between the PM and mitochondria, which are less extensive [48], have received less attention. In HeLa cells co-expressing the PM β_2_-adrenoceptor tagged with mCh (β_2_AR-mCh), TOM20-GFP-FRB and RNb-FKBP, rapamycin caused rapid recruitment of β_2_AR-mCh within the PM to mitochondria at discrete puncta that grew larger with time (Fig. 12A-E and ***Additional file 8: Video 7***). Recruitment was not seen in the absence of co-expressed RNb-FKBP (Fig. 12F). Rapamycin also triggered similar punctate accumulation of β_2_AR at mitochondria in COS-7 cells expressing β_2_AR-GFP, TOM20-mCh-FRB and GNb-FKBP (***Additional file 1: Fig. S4***). In similar analyses of ER-mitochondria and PM-mitochondria MCS, the initial punctate colocalization of proteins was shown to report native MCS, which grew larger with time as rapamycin zipped the proteins together [48]. Our results are consistent with that interpretation. In most cases, β_2_AR were recruited to only one or two discrete sites on each mitochondrion, which expanded during prolonged incubation with rapamycin, but without the appearance of new sites (Fig. 12D and E, and ***Additional file 1: Fig. S4***). Rapamycin had no evident effect on recruiting new mitochondria to the PM, but it did cause accumulation of tagged TOM70 at MCS and depletion of TOM70 from the rest of each mitochondrion, indicating mobility of TOM70 within the OMM (***Additional file 1: Fig. S4***). Our results suggest that inducible cross-linking using RNb-FKBP or GNb-FKBP identifies native MCS between mitochondria and PM, with each mitochondrion forming only one or two MCS with the PM. We have not explored the functional consequences of these restricted MCS, but we speculate that they may identify sites where proteins involved in communication between the PM and mitochondria are concentrated, facilitating, for example, phospholipid transfer [49], the generation of ATP microdomains [50], or Ca^2+^ exchanges between mitochondria and store-operated Ca^2+^ entry (SOCE) [51] or PM Ca^2+^-ATPases [52].

**Fig 12.**
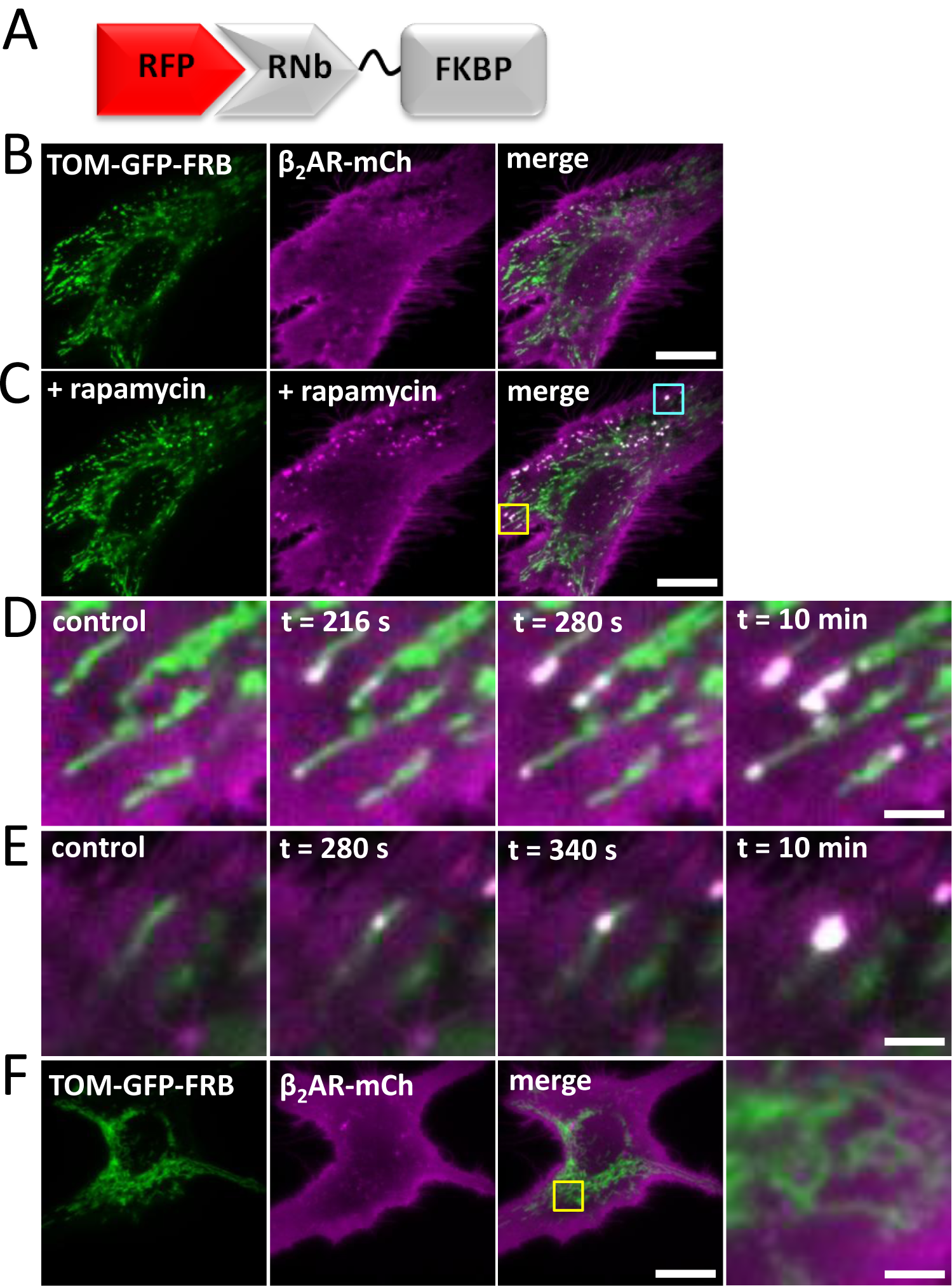
Recruitment of proteins to native PM-mitochondria MCS using RNb-FKBP. (**A**) Schematic of RNb-FKBP fusion bound to RFP. (**B, C**) HeLa cells co-expressing RNb-FKBP, mitochondrial TOM70-GFP-FRB and β_2_AR-mCh were imaged using TIRFM before (B) and after (C) treatment with rapamycin (100 nM, 10 min). Scale bar 10 µm. (**D, E**) Enlarged images from C of the yellow box (D) and cyan box (E) show punctate recruitment of β_2_AR-mCh to individual mitochondria at the indicated times after addition of rapamycin. Scale bars 1.25 µm. (**F**) TIRFM images of HeLa cells co-expressing mitochondrial TOM70-GFP-FRB and β_2_AR-mCh in the presence of rapamycin (100 nM, 10 min) show no recruitment in the absence of co-expressed RNb-FKBP. The yellow box shows a region enlarged in the subsequent image. Scale bars 10 µm (main images) and 2.5 μm (enlargement). Results (B-F) are representative of 5 independent experiments.

We next tested whether PM proteins could be recruited to the MCS between ER-PM that are important for SOCE and lipid transfer [53]. In response to rapamycin, mCh-Orai1, the PM Ca^2+^ channel that mediates SOCE [54], was recruited by RNb-FKBP to ER-PM MCS labelled with the marker GFP-MAPPER-FRB [55] (Fig. 13A and B). Recruitment was not observed in the absence of RNb-FKBP (Fig. 13C). We conclude that the method identifies native ER-PM MCS during the initial phase of Nb recruitment, and the Nb subsequently exaggerates these MCS.

**Fig 13.**
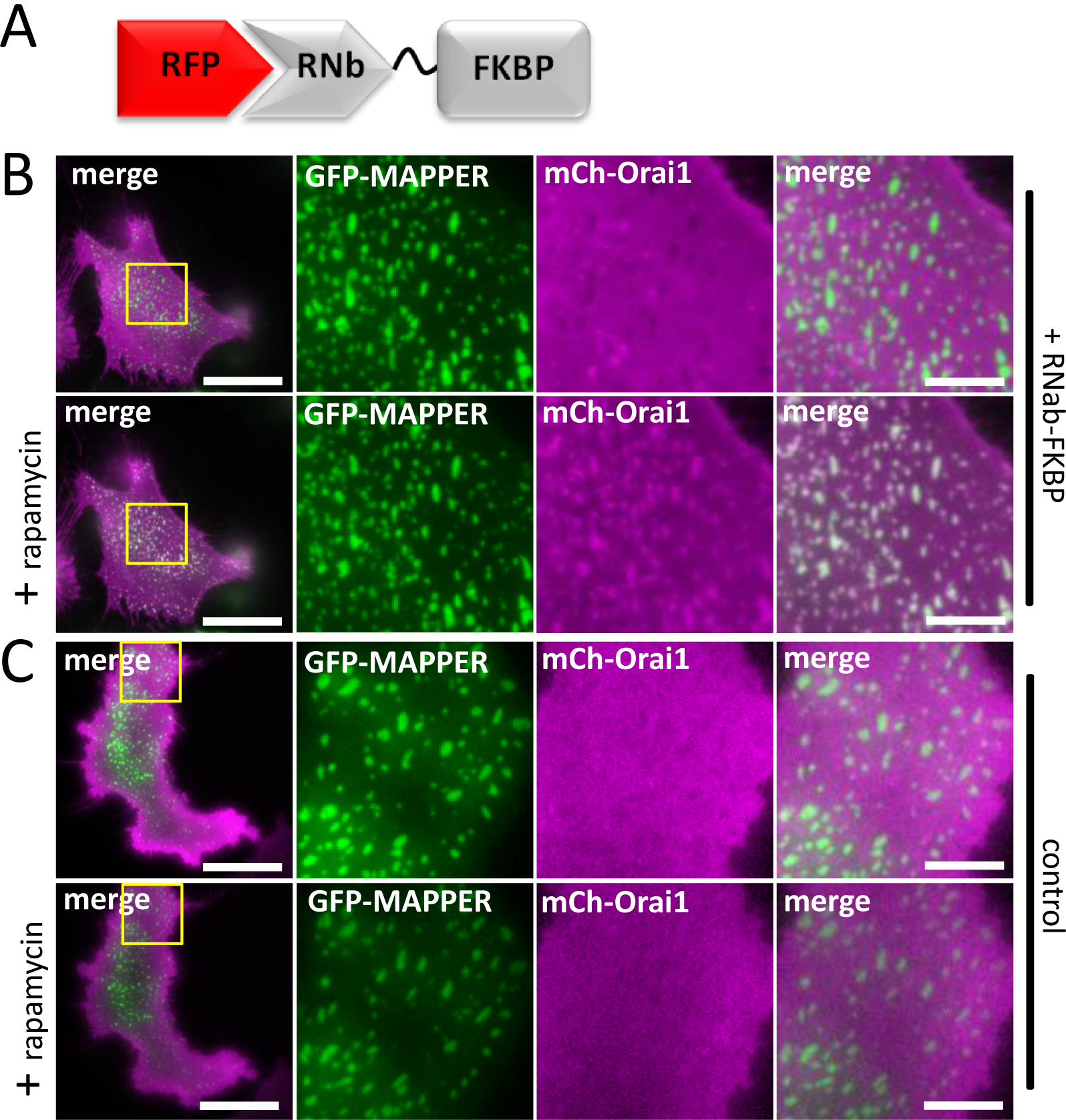
Recruitment of PM proteins to ER-PM MCS using RNb-FKBP. (**A**) Schematic of RNb-FKBP fusion bound to RFP. (**B**) HeLa cells co-expressing RNb-FKBP, mCh-Orai1 and the ER-PM junction marker GFP-MAPPER-FRB were imaged using TIRFM. A representative cell (n = 5) is shown before (top row) and after (bottom row) treatment with rapamycin (100 nM, 10 min). The boxed region is shown enlarged in subsequent images. (**C**) HeLa cells co-expressing mCh-Orai1 and GFP-MAPPER-FRB alone were imaged using TIRFM. A representative cell (n = 3) is shown before (top row) and after (bottom row) treatment with rapamycin (100 nM, 10 min). The boxed region is shown enlarged in subsequent images. The results show no recruitment in the absence of co-expressed RNb-FKBP. Scale bars (B, C) 10 µm (main images) and 2.5 µm (enlargements).

One of the least explored MCS is that between lysosomes and mitochondria [56]. Recent evidence shows that these MCS control the morphology of both organelles [57] and probably mediate exchange of cholesterol and other metabolites between them [58]. We assessed whether the nanobody fusions could be used to inducibly recruit lysosomes to mitochondria. GNb-FKBP enabled recruitment of lysosomes labelled with LAMP1-GFP to mitochondria labelled with TOM20-mCh-FRB, in response to rapamycin (Fig. 14A-C). Lysosomes were not recruited to mitochondria in the absence of GNb-FKBP (Fig. 14D).

**Fig 14.**
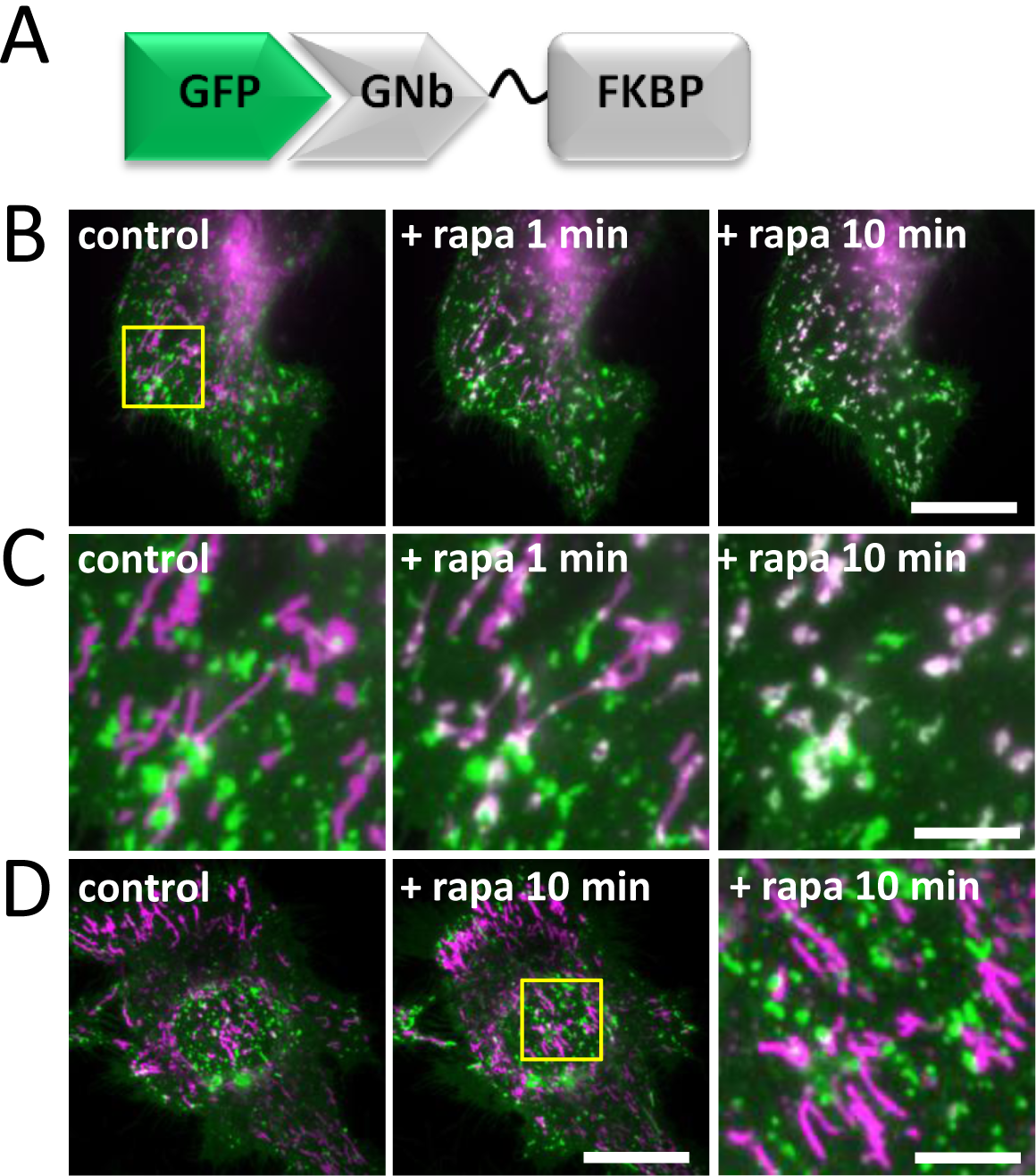
Inducible recruitment of lysosomes to mitochondria using GNb-FKBP. (**A**) Schematic of GNb-FKBP fusion bound to GFP. (**B**) HeLa cells co-expressing mitochondrial TOM70-mCh-FRB (magenta), lysosomal LAMP1-GFP (green) and GNb-FKBP were imaged using TIRFM. Merged images of a representative cell (n = 5) are shown before and at times after treatment with rapamycin (rapa, 100 nM). Scale bar 10 µm. (**C**) Enlargements of the boxed region in (B). Scale bar 2.5 μm. (**D**) HeLa cells co-expressing TOM70-mCh-FRB (magenta) and lysosomal LAMP1-GFP (green) were imaged using TIRFM. A representative cell (n = 3) is shown before and after treatment with rapamycin (100 µm, 10 min); there is no recruitment in the absence of co-expressed GNb-FKBP. The yellow box shows a region enlarged in the subsequent image. Scale bars 10 µm (main images) and 2.5 μm (enlargement).

### Cross-linking RFP-tagged and GFP-tagged proteins

We generated a dimeric nanobody (GNb-RNb) that binds simultaneously to GFP and RFP (Fig. 15A), and demonstrated its utility by crosslinking a variety of GFP-tagged and RFP-tagged proteins. Cytosolic GFP, normally diffusely distributed in the cytosol (data not shown), was recruited to nuclei by H2B-mCh (Fig. 15B) or to mitochondria by TOM20-mCh (Fig. 15C). In the presence of GNb-RNb, mCh-Orai1 and endogenously tagged GFP-IP_3_R1 formed large co-clusters (Fig 15D) that differed markedly from the distributions of GFP-IP_3_R1 (Fig. 10F) and mCh-Orai1 (Fig 13) in the absence of crosslinking. Consistent with earlier results (Fig. 12 and ***Additional file 1: Fig. S4***), β_2_AR-mCh, which is normally diffusely distributed in the PM, formed mitochondria-associated puncta when crosslinked to mitochondria expressing TOM20-GFP (Fig. 15E). Whole organelles could also be crosslinked. Co-expression of LAMP1-GFP and LAMP1-mCh labelled small, mobile lysosomes in control cells (Fig. 15F), while additional co-expression of GNb-RNb caused accumulation of lysosomes into large clusters (Fig. 15G).

**Fig 15.**
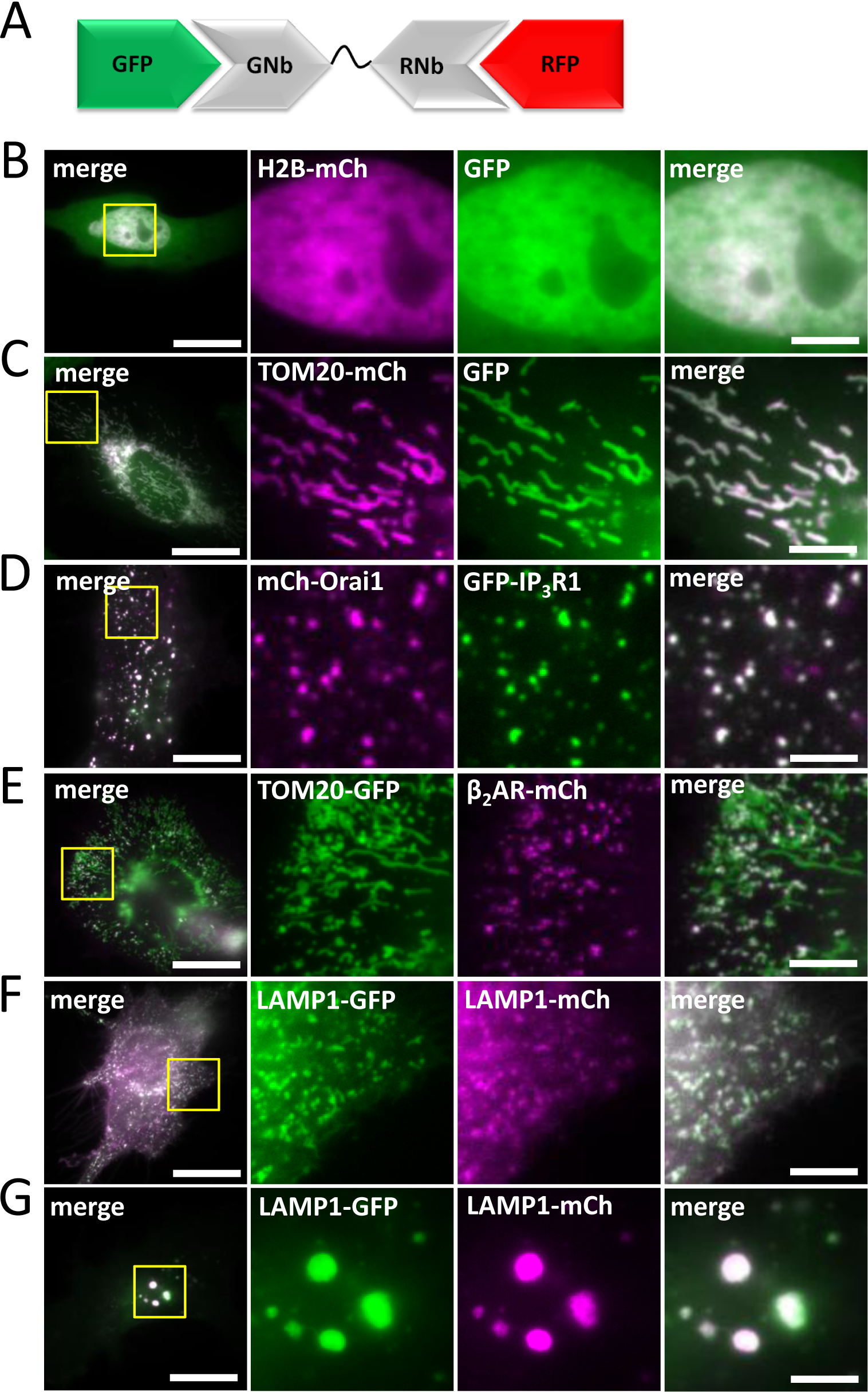
Crosslinking GFP-tagged and RFP-tagged proteins and organelles using GNb-RNb. (**A**) Schematic of GNb-RNb bound to GFP and RFP. (**B-E**) HeLa cells co-expressing the tagged proteins indicated with GNb-RNb were imaged using epifluorescence microscopy (B) or TIRFM (C-E). Representative cells (n = 5-7) are shown. Control images for GFP-IP_3_R1 are shown in Fig. 10 and ***Additional file 1: Fig. S3***. (**F, G**) HeLa cells co-expressing LAMP1-GFP and LAMP1-mCh in the absence (F) or presence (G) of co-expressed GNb-RNb were imaged using TIRFM. Representative cells (n = 5) are shown. Scale bars (B-G) 10 µm (main images) and 2.5 µm (enlargements of boxed areas).

This crosslinking of GFP and RFP was made rapidly inducible with an RNb-FRB fusion that hetero-dimerizes with GNb-FKBP in the presence of rapamycin (Fig. 16A). Co-expression of GNb-FKBP with RNb-FRB in cells co-expressing TOM20-GFP and mCh-Sec61β led to rapid colocalization of GFP and mCh after addition of rapamycin (Fig. 16B and C, and ***Additional file 9: Video 8***). Similar results were obtained with RNb-FKBP and GNb-FRB (***Additional file 1: Fig. S5***). We conclude that GNb-FKBP and RNb-FRB provide a rapidly inducible system for crosslinking any GFP-tagged protein to any RFP-tagged protein.

**Fig 16.**
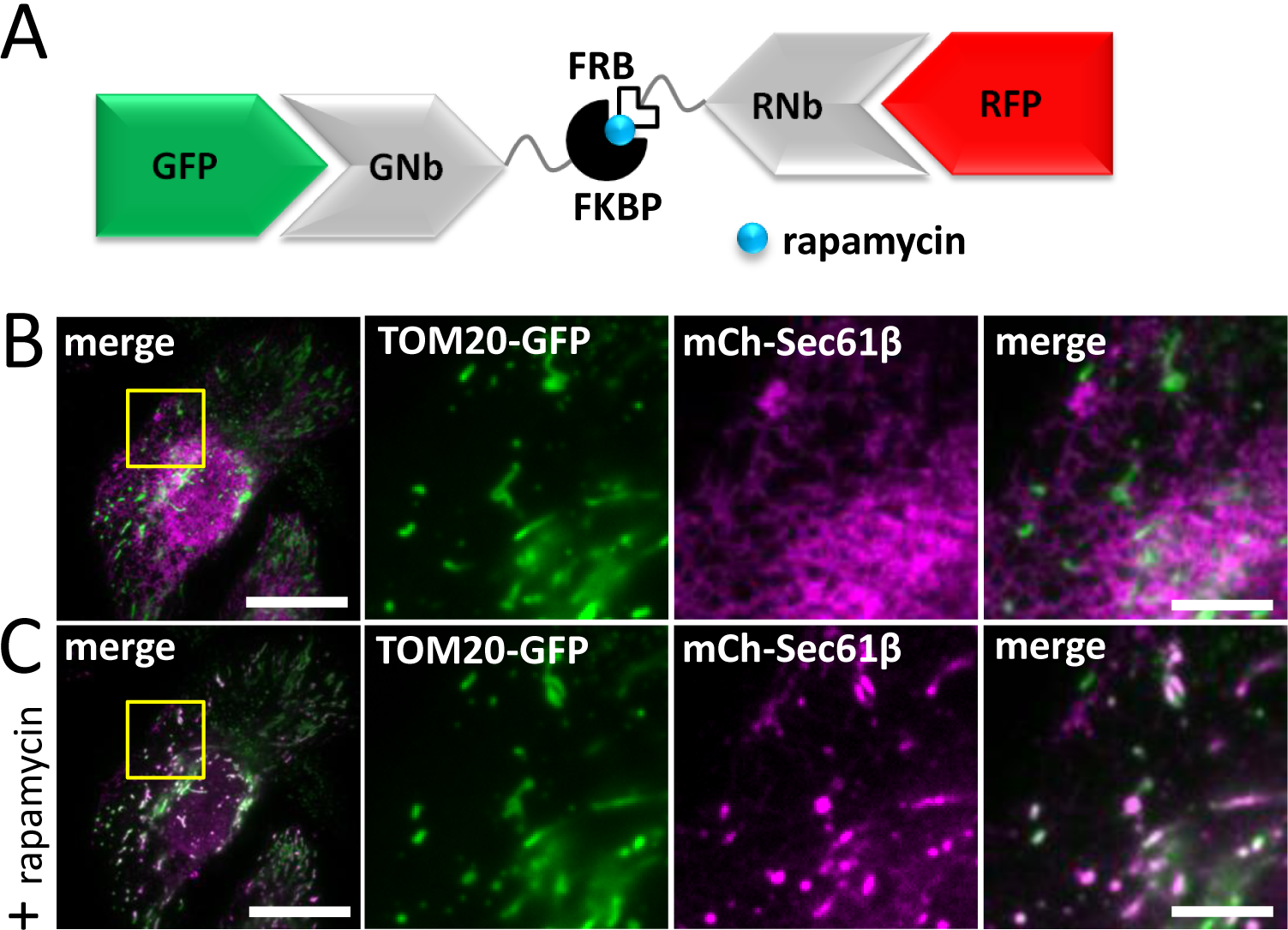
Inducible crosslinking of RFP-tagged and GFP-tagged proteins with GNb-FKBP and RNb-FRB. (**A**) Schematic of the nanobody fusions used, with rapamycin shown as a blue sphere. (**B**, **C**) HeLa cells co-expressing GNb-FKBP, RNb-FRB, TOM20-GFP and mCh-Sec61β were imaged using TIRFM. A representative cell (n = 3) is shown before (B) and after (C) treatment with rapamycin (100 nM, 10 min). Scale bars 10 µm (main images) and 2.5 µm (enlargements of boxed areas).

### Targeting secretory compartments with lumenal nanobodies

GNb and RNb were directed to the lumen of the secretory pathway by addition of an N-terminal signal sequence, giving ssGNb and ssRNb. Targeting of ssGNb-mCh to the Golgi, ER network,or ER-PM MCS was achieved by co-expression of organelle markers with lumenal FP tags (Fig. 17A and B). In each case, there was significant colocalization of green and red proteins. Similar targeting of ssRNb-GFP to the ER network or ER-PM MCS was achieved by co-expression with mCh-tagged lumenal markers of these organelles (Fig. 17C and D). These results demonstrate that ssGNb and ssRNb fusions can be directed to the lumen of specific compartments of the secretory pathway.

**Fig 17.**
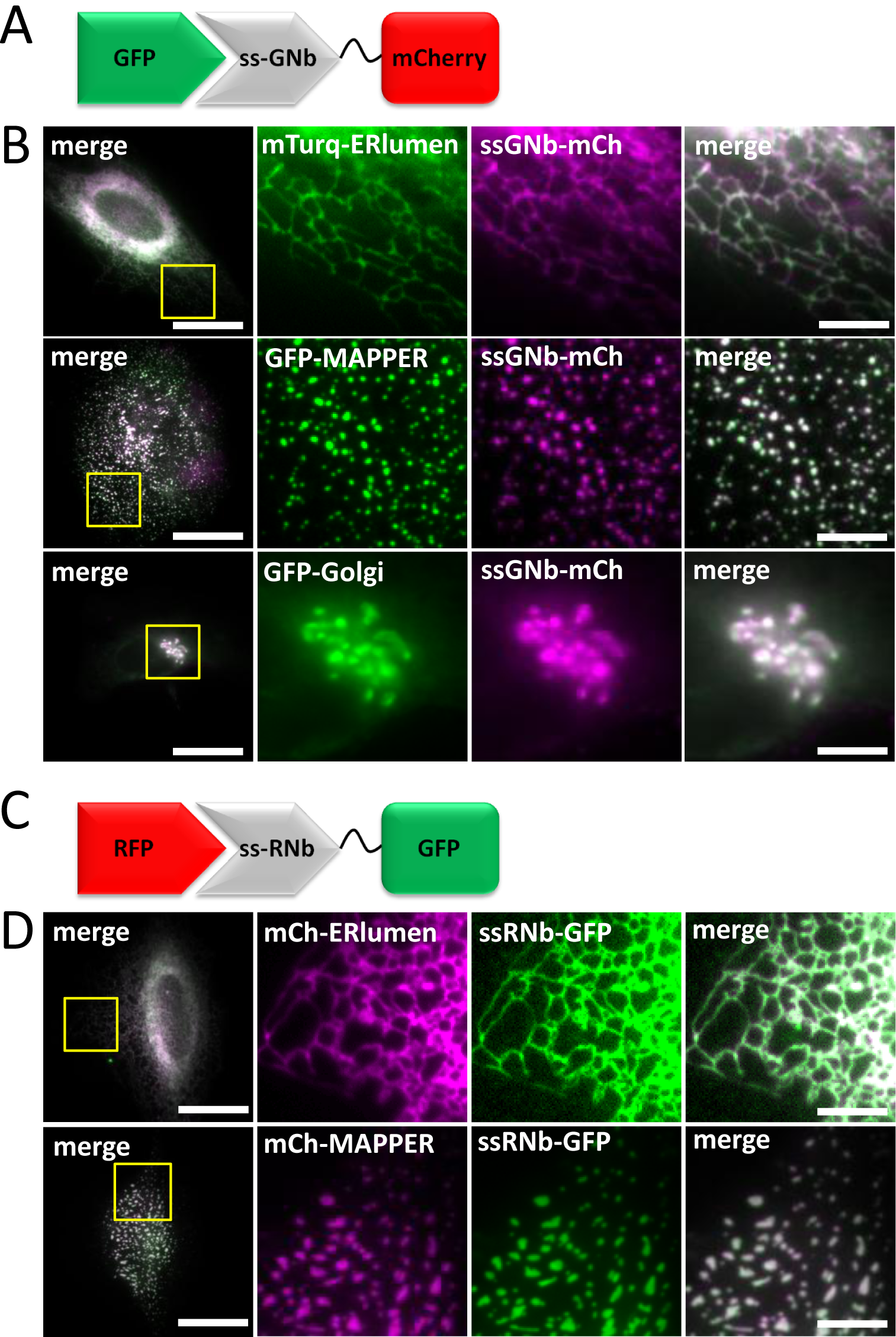
Nanobody fusions can be targeted to different lumenal compartments of the secretory pathway. (**A**) Schematic of ssGNb-mCh bound to GFP. (**B**) HeLa cells co-expressing ssGNb-mCh and either the lumenal ER marker mTurquoise2-ERlumen, the marker of ER-PM junctions GFP-MAPPER, or the Golgi marker GFP-Golgi. Cells were imaged using epifluorescence microscopy. Representative cells are shown. Colocalization values were: mTurquoise2-ERlumen (*r* = 0.96 ± 0.03, n = 10); GFP-MAPPER (*r* = 0.94 ± 0.02, n = 5); and GFP-Golgi (*r* = 0.91 ± 0.06, n = 4). (**C**) Schematic of ssRNb-GFP bound to RFP. (**D**) HeLa cells co-expressing ssRNb-GFP and either mCh-ERlumen or mCh-MAPPER were imaged using epifluorescence microscopy. Representative cells are shown. Colocalization values were: mCh-ERlumen (*r* = 0.98 ± 0.009, n = 9) and mCh-MAPPER (*r* = 0.93 ± 0.07, n = 13. Scale bars 10 µm (main images) and 2.5 µm (enlargements of boxed regions).

Fluorescent Ca^2+^ sensors targeted to the lumen of the entire ER [59, 60] are widely used and have considerably advanced our understanding of Ca^2+^ signalling [61, 62]. Fluorescent Ca^2+^ sensors targeted to ER sub-compartments and the secretory pathway have received less attention but have, for example, been described for the Golgi [63, 64]. Our nanobody methods suggest a generic approach for selective targeting of lumenal Ca^2+^ indicators. Fusion of ssRNb to GCEPIA1 or GEM-CEPIA [60] provided ssRNb-GCEPIA1 and ssRNb-GEMCEPIA (Fig. 18A). These fusions were targeted to the lumenal aspect of ER-PM junctions by co-expression with mCh-MAPPER [7] (Fig. 18C and D). Fusion of ssGNb to the low-affinity Ca^2+^ sensors LAR-GECO1 [59] or RCEPIA1 [60] provided ssGNb-LARGECO1 and ssGNb-RCEPIA1 (Fig. 18B). These fusions allowed targeting to ER-PM junctions labelled with GFP-MAPPER (Fig. 18E and F). The targeted Ca^2+^ sensors responded appropriately to emptying of intracellular Ca^2+^ stores by addition of ionomycin in Ca^2+^-free medium (Fig. 18G-K). These results confirm that Ca^2+^ sensors targeted to a physiologically important ER sub-compartment, the ER-PM junctions where SOCE occurs, report changes in lumenal [Ca^2+^]. Our results demonstrate that nanobody fusions can be targeted to lumenal sub-compartments of the secretory pathway and they can report [Ca^2+^] within physiologically important components of the ER.

**Fig 18.**
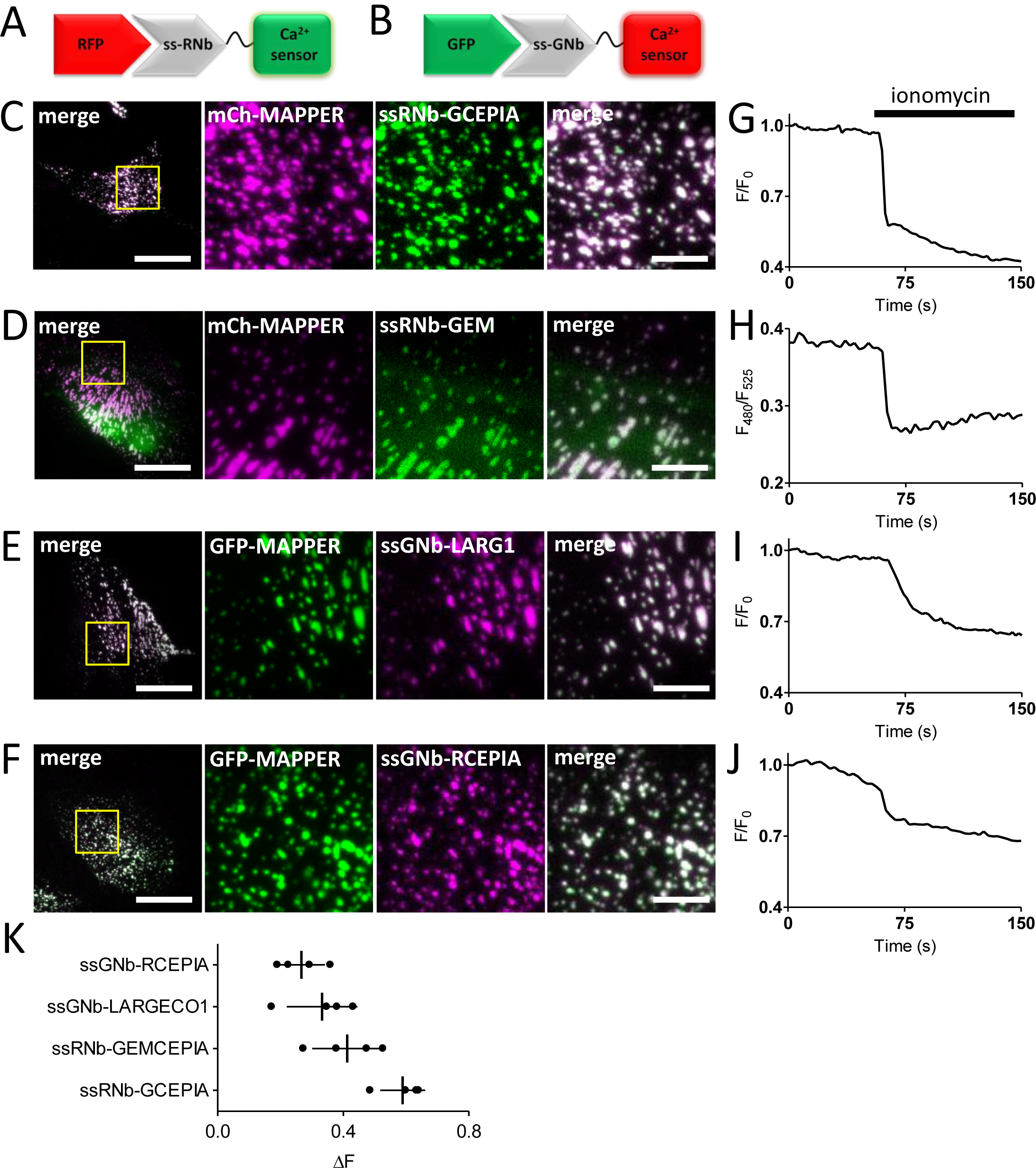
Nanobody-mediated targeting of low-affinity Ca^2+^ sensors allows measurement of changes in [Ca^2+^] in an ER sub-compartment at ER-PM MCS. (**A**) Schematic of ssRNb-Ca^2+^ sensor bound to RFP. (**B**) Schematic of ssGNb-Ca^2+^ sensor bound to GFP. (**C-F**) HeLa cells co-expressing the indicated combinations of mCh-MAPPER, GFP-MAPPER, ssRNb-GCEPIA (ssRNb-GC), ssRNb-GEMCEPIA (ssRNb-GEM; image is shown for the 525-nm emission channel), ssGNb-LAR-GECO1 (ssGNb-LGECO) or ssGNb-RCEPIA were imaged in Ca^2+^-free HBS using TIRFM. Yellow boxes indicate regions enlarged in subsequent images. Scale bars 10 µm (main images) and 2.5 µm (enlargements). (**G-J**) Timecourses of fluorescence changes recorded from cells co-expressing mCh-MAPPER and ssRNb-GCEPIA (G), mCh-MAPPER and ssRNb-GEMCEPIA (H), GFP-MAPPER and ssGNb-LAR-GECO1 (ssGNb-LARG1) (I) and GFP-MAPPER and ssGNb-RCEPIA (J) in response to emptying of intracellular Ca^2+^ stores with ionomycin (5 µM). (**K**) Summary results (with mean ± SD, n = 4 cells) show fractional decreases (ΔF) in either fluorescence or emission ratio (for ssRNb-GEM) recorded 90 s after addition of ionomycin,

## Discussion

The spatial organization of the cell interior influences all cellular activities and it is a recurrent theme in intracellular signalling [65, 66]. Hence, tools that can visualize and manipulate the spatial organization of intracellular components are likely to find widespread application. We introduce a toolkit of plasmids encoding functionalized nanobodies against common FP tags, including CFP, GFP, YFP and RFPs (Fig. 1). Use of this toolkit is supported by genome-wide collections of plasmids, cells and organisms expressing proteins tagged with GFP and RFP [10-17, 19], and by facile methods for heterologous expression of tagged proteins or editing of endogenous genes to encode FP tags [5, 6]. The functionalized nanobodies provide new approaches to studying intracellular signalling in live cells.

Our toolkit expands the repertoire of functionalized RFP-binding nanobodies, which are less developed than their GFP-binding counterparts [67]. The RNb fusions provide new opportunities to use RFP, which often has advantages over GFP. For example, RFP is spectrally independent from blue-green sensors, which are usually superior to their red counterparts [30, 32]; from the CALI probe, fluorescein; and from optogenetic modules, which are often operated by blue-green light [68].

Nanobody-sensor fusions allow targeting of sensors to specific proteins and organelles (Figs. 2-6), and will aid visualization of signalling within cellular microdomains. Fusion of nanobodies to the Ca^2+^ sensors G-GECO1.2, R-GECO1.2 and LAR-GECO1.2 [30] (Figs. 3 and 4), which have relatively low affinities for Ca^2+^ (K_D_ values of 1.2 µM, 1.2 µM and 10 µM, respectively), should facilitate selective detection of the relatively large, local rises in [Ca^2+^]_c_ that are important for cell signalling [27]. The GEM-GECO Ca^2+^ sensor [30], H^+^ sensors [31, 32] and ATP/ADP sensors [33] used for nanobody fusions are poised to detect fluctuations of their ligands around resting concentrations in the cell (Figs. 4-6).

Relative to direct fusions of sensors to proteins of interest, nanobody-sensor fusions have several advantages. Firstly, the generic nanobody toolkit (Fig. 1) can be combined with collections of FP-tagged proteins to provide many combinations; each would otherwise require expression of a unique construct that may or may not function normally. Secondly, each sensor is attached to the same entity (nanobody), which binds to the same partner (FP). Since the biophysical and biochemical properties of sensors may be influenced by their fusion partners, this provides greater confidence that sensors despatched to different locations will respond similarly to their analyte.

Nanobodies allow re-colouring of FPs with alternative fluorophores that may have advantageous properties. For example, recolouring of RFP-tagged proteins with RNb-GFP (Fig. 2B) enables visualization of organelles with GFP, which has enhanced photophysical properties relative to RFPs. Nanobody-SNAPf fusions can be used to attach fluorescent dyes, including CALI probes and far-red fluorophores, to FP tags (Figs. 7 and 8). Longer excitation wavelengths cause less phototoxicity and allow greater penetration through tissue, which may be useful in studies of transgenic organisms and tissues. We also envisage live-cell applications in pulse-chase analyses and using super-resolution microscopy, Förster resonance energy transfer (FRET) and fluorescence lifetime imaging.

Membrane-permeant forms of the SNAP ligand, O^6^-benzylguanine, are available conjugated to conventional Ca^2+^ indicators (Fura-2FF, Indo-1 and BOCA-1), which are brighter than genetically-encoded indicators [69-71]; to derivatives of the two-photon fluorophore naphthalimide [72]; to the hydrogen peroxide sensor nitrobenzoylcarbonylfluorescein [73]; and to reversible chemical dimerizers [74, 75]. Nanobody-SNAPf fusions will allow facile targeting of these modules to any protein or organelle tagged with RFP or GFP.

Cross-linking methods have many applications in cell biology, including stabilizing protein interactions (eg, for pull-downs), identifying and manipulating MCS, enforcing protein interactions (eg, receptor dimerization), redirecting proteins to different subcellular locations (eg, knocksideways) and many more. Functionalized nanobodies provide many additional opportunities to regulate protein associations. The nanobody-FKBP/FRB fusions, for example, allow rapid rapamycin-mediated crosslinking of any pair of proteins tagged with GFP/RFP, or tagged with either FP and any of the many proteins already tagged with FKBP or FRB [76] (Figs. 10 and 12-16). Nanobody-FKBP fusions may allow crosslinking to SNAP-tagged proteins [75], and the nanobody-SNAPf fusions to HaloTag-tagged proteins [74] and FKBP-tagged proteins [75]. RNb-zdk1 fusions allow photo-inducible crosslinking of RFP-tagged proteins to LOV-tagged proteins [46] (Fig. 11). Nanobodies that crosslink GFP-tagged proteins to RFP-tagged proteins (GNb-RNb; and the GNb-FKBP/RNb-FRB and GNb-FRB/RNb-FKBP pairings) may have the most applications, as they can take fullest advantage of the numerous combinations of existing RFP and GFP-tagged proteins (Figs. 15 and 16).

Functionalized nanobodies directed to lumenal compartments of the secretory pathway would provide useful tools, but they are under-developed. Their potential is shown by nanobodies retained within the ER, which restrict onward trafficking of target proteins and inhibit their function [77]. We show that functionalized nanobodies, including nanobody-Ca^2+^ sensors, can be directed to sub-compartments of the secretory pathway (Figs. 17 and 18). Lumenal Ca^2+^ provides a reservoir within the ER, Golgi and lysosomes that can be released by physiological stimuli to generate cytosolic Ca^2+^ signals [78, 79]. Compartmentalization of Ca^2+^ stores within the ER [63] and Golgi [79] adds to the complexity of lumenal Ca^2+^ distribution in cells. Furthermore, lumenal Ca^2+^ itself regulates diverse aspects of cell biology, including SOCE [54], sorting of cargo in the Golgi [80], binding of ERGIC-53 to cargoes within the ER-Golgi intermediate compartment (ERGIC) [81], and exocytosis of neurotransmitters by secretory vesicles [82, 83]. Hence, there is a need for tools that can effectively report lumenal [Ca^2+^] within this complex lumenal environment. The lumenal nanobody-Ca^2+^ sensors detected changes in lumenal [Ca^2+^] at the ER-PM MCS where SOCE occurs (Fig. 18).

In addition to nanobodies, other protein-based binders, including single-domain antibodies, designed ankyrin-repeat proteins (DARPINs), affimers, anticalins, affibodies and monobodies have been developed to recognise many important intracellular proteins [2, 84-86]. These binding proteins can be easily transplanted into the fusion scaffolds described to maximize their exploitation.

## Conclusions

We present a toolkit of plasmids encoding functionalized nanobodies directed against common fluorescent protein tags, which will allow a wide range of applications and new approaches to studying intracellular signalling in live cells. We illustrate some applications and demonstrate, for example, that IP_3_ receptors deliver Ca^2+^ to the OMM of only some mitochondria, and that MCS between mitochondria and the plasma membrane occur at only one or two sites on each mitochondrion.

## Materials and Methods

### Materials

Human fibronectin was from Merck Millipore. Ionomycin was from Apollo Scientific (Stockport, UK). Rapamycin was from Cambridge Bioscience (Cambridge, UK). SNAP substrates were from New England Biolabs (Hitchin, UK). Other reagents, including histamine and nigericin, were from Sigma-Aldrich.

### Plasmids

Sources of plasmids encoding the following proteins were: mCherry-C1 (Clontech #632524); mCherry-N1 (Clontech #632523); EGFP-N1 (Clontech #6085-1); GFP-ERcyt, mCherry-ERcyt and mTurquoise2-ERcyt (GFP, mCherry or mTurquoise2 targeted to the cytosolic side of the ER membrane via the ER-targeting sequence of the yeast UBC6 protein) [87]; mCherry-ERlumen (Addgene #55041, provided by Michael Davidson); LAMP1-mCherry [88]; TPC2-mRFP [89]; TOM20-mCherry (Addgene #55146, provided by Michael Davidson); CIB1-mRFP-MP (Addgene #58367) [44]; CIB1-mCerulean-MP (Addgene #58366) [44]; H2B-GFP (Addgene #11680) [90]; TOM20-LOV2 (Addgene #81009) [46]; mCherry-Sec61β [91]; GFP-MAPPER [55]; GFP-CaM (Addgene #47602, provided by Emanuel Strehler); TOM70-mCherry-FRB (pMito-mCherry-FRB, Addgene #59352) [92]; pmTurquoise2-Golgi (Addgene #36205) [93]; pTriEx-mCherry-zdk1 (Addgene #81057) [46]; pTriEx-NTOM20-LOV2 (Addgene #81009) [46]; β_2_AR-mCFP (Addgene #38260) [94]; pCMV-G-CEPIA1er (Addgene #58215) [60]; pCMV-R-CEPIA1er (Addgene #58216) [60]; pCIS-GEM-CEPIA1er (Addgene #58217) [60]; CMV-ER-LAR-GECO1 and CMV-mito-LAR-GECO1.2 [59]; mCherry-MAPPER and mCherry-Orai1 [7].

H2B-mCh was made by transferring H2B from H2B-GFP to pmCherry-N1 (Clontech) using *Kpn*I/*Bam*HI. LAMP1-GFP was made by transferring LAMP1 from LAMP1-mCherry into pEGFP-N1 (Clontech) using *Eco*RI/*Bam*HI. β_2_AR-mCherry was made by transferring β_2_AR from β_2_AR-mCFP to pmCherry-N1 (Clontech) using *Nhe*I/*Xho*I. β_2_AR-GFP was made by transferring GFP from pEGFP-N1 (Clontech) into β_2_AR-mCherry using *Xho*I/*Not*I. The mCherry-Golgi plasmid was made by transferring mCherry from pmCherry-N1 into pEYFP-Golgi (Clontech) using *Age*I/*Not*I. GFP-Golgi was made by transferring GFP from pEGFP-N1 (Clontech) into Golgi-mCherry using *Age*I/*Not*I. TOM20-GFP was made by transferring EGFP from pEGFP-N1 into TOM20-mCherry using *Bam*HI/*Not*I. TOM70-GFP-FRB was made by insertion of EGFP from pEGFP-N1 into TOM70-mCh-FRB using *Age*I/*Bsr*GI. SNAPf-pcDNA3.1(+) was made by transferring SNAPf from pSNAPf (New England Biolabs) to pcDNA3.1 (+) using *Nhe*I/*Not*I.

DNA constructs encoding GNb and RNb were synthesized as DNA Strings (ThermoFisher) and introduced by Gibson assembly (Gibson Assembly Master Mix, New England Biolabs) into pcDNA3.1(+) digested with *Bam*HI/*Eco*RI. Sequences encoding GNb and RNb are shown in ***Additional file 1: Fig. S6.*** Plasmids encoding nanobody fusion constructs (Fig. 1) were constructed from the GNb and RNb plasmids using PCR, restriction digestion and ligation, or synthetic DNA Strings and Gibson assembly, and their sequences were confirmed.

GNb-mCherry was made by PCR of pmCherry-N1 using forward (ACTGGATCCATGGTGAGCAAGGGCGAG) and reverse (GTACTCGAGCTACTTGTACAGCTCGTCCATGC) primers, followed by insertion into GNb-pcDNA3.1(+) using *Bam*HI/*Xho*I. RNb-GFP was made by PCR of pEGFP-N1 using forward (ACTGGATCCATGGTGAGCAAGGGCGAG) and reverse (GTACTCGAGCTACTTGTACAGCTCGTCCATGC) primers, followed by insertion into RNb-pcDNA3.1(+) using *Bam*HI/*Xho*I. RNb-mCerulean-MP was made by PCR of RNb using forward (ATGCTAGCAAGCTTGCCACCATGGCTC) and reverse (ATACCGGTGAGGATCCAGAGCCTCCGC) primers, followed by insertion into CIB1-mCerulean-MP using *Nhe*I/*Age*I. GNb-mRFP-MP was made by PCR of GNb-FKBP with forward (TAGCTAGCGCCACCATGGCTCAGGTG) and reverse (CGACCGGTACGGACACGGTCACTTGGG) primers, and insertion into CIB1-mRFP1-MP using *Nhe*I/*Age*I. GNb-SNAPf and RNb-SNAPf were made by PCR of GNb-pcDNA3.1(+) and RNb-pCDNA3.1(+) using forward (CAGCTAGCTTGGTACCGAGCTCAAGCTTGC) and reverse (ATGAATTCAGATCCCCCTCCGCCAC) primers, followed by insertion into SNAPf-pcDNA3.1 (+) using *Nhe*I/*Eco*RI. GNb-LAR-GECO1.2 was made by PCR of CMV-mito-LAR-GECO1.2 using forward (CAGGATCCATGGTCGACTCTTCACGTCGTAAGTGG) and reverse (GTACTCGAGCTACTTCGCTGTCATCATTTGTACAAACT) primers, followed by insertion into GNb-pcDNA3.1(+) using *Bam*HI/*Xho*I. RNb-GGECO1.2 was made by PCR of CMV-G-GECO1.2 using forward (CAGGATCCATGGTCGACTCATCACGTCGTAAG) and reverse (TACGATGGGCCCCTACTTCGCTGTCATCATTTGTACAAACTCTTC) primers, followed by insertion into RNb-pcDNA3.1(+) using *Bam*HI/*Apa*I. RNb-Perceval-HR was made by PCR of Perceval-HR with forward (AAGCGGCCGCTATGAAAAAGGTTGAATCCATCATCAGGCC) and reverse (ATCTCGAGTCACAGTGCTTCCTTGCCCTC) primers, followed by insertion into RNb-pcDNA3.1(+) using *Not*I/*Xho*I.

ss-GNb-mCherry was made by inserting mCherry from GNb-mCherry into ss-GNb-FKBP using *Bam*HI/*Not*I. ss-RNb-GFP was made by inserting GFP from RNb-GFP into ss-RNb-pcDNA3.1(+) using *Bam*HI/*Not*I. ss-GNb-RCEPIA was made by transferring RCEPIA from pCMV-R-CEPIA1er to ss-RNb-pcDNA3.1(+) using *Bam*HI/*Not*I. ss-GNb-LAR-GECO1 was made by transferring a DNA String encoding ss-GNb into CMV-ER-LAR-GECO1 using *Hind*III/*Bam*HI. ss-RNb-GCEPIA was made by transferring GCEPIA from pCMV-G-CEPIA1er to ss-RNb-pcDNA3.1(+) using *Bam*HI/*Not*I. ss-RNb-GEMCEPIA was made by transferring GEM-CEPIA from pCIS-GEM-CEPIA1er to ss-RNb-pcDNA3.1(+) using *Bam*HI/*Not*I.

### Cell culture and transient transfection

HeLa and COS-7 cells (American Type Culture Collection) were cultured in Dulbecco’s modified Eagle’s medium/F-12 with GlutaMAX (ThermoFisher) supplemented with foetal bovine serum (FBS, 10%, Sigma). Cells were maintained at 37°C in humidified air with 5% CO_2_, and passaged every 3-4 days using Gibco TrypLE Express (ThermoFisher). For imaging, cells were grown on 35-mm glass-bottomed dishes (#P35G-1.0-14-C, MatTek) coated with human fibronectin (10 µg.ml^−1^). Cells were transfected, according to the manufacturer’s instructions, using TransIT-LT1 (GeneFlow) (1 µg DNA per 2.5 µl reagent). Short tandem repeat profiling (Eurofins, Germany) was used to authenticate the identity of HeLa cells [7]. Screening confirmed that all cells were free of mycoplasma infection.

### Fluorescence microscopy and analysis

Cells were washed prior to imaging at 20°C in HEPES-buffered saline (HBS: NaCl 135 mM, KCl 5.9 mM, MgCl_2_ 1.2 mM, CaCl_2_ 1.5 mM, HEPES 11.6 mM, D-glucose 11.5 mM, pH 7.3). Ca^2+^-free HBS lacked CaCl_2_ and contained EGTA (1 mM). For manipulations of intracellular pH, cells were imaged in modified HBS (MHBS: KCl 140 mM, MgCl_2_ 1.2 mM, CaCl_2_ 1.5 mM, HEPES 11.6 mM, D-glucose 11.5 mM, pH 7.2). The H^+^/K^+^ ionophore nigericin (10 μM) was added 5 min before imaging to equilibrate intracellular and extracellular pH, and the extracellular pH was then varied during imaging by exchanging the MHBS (pH 6.5 or pH 8).

Fluorescence microscopy was performed at 20°C as described previously [7] using an inverted Olympus IX83 microscope equipped with a 100× oil-immersion TIRF objective (numerical aperture, NA 1.49), a multi-line laser bank (425, 488, 561 and 647 nm) and an iLas2 targeted laser illumination system (Cairn, Faversham, Kent, UK). Excitation light was transmitted through either a quad dichroic beam splitter (TRF89902-QUAD) or a dichroic mirror (for 425 nm; ZT442rdc-UF2) (Chroma). Emitted light was passed through appropriate filters (Cairn Optospin; peak/bandwidth: 480/40, 525/50, 630/75 and 700/75 nm) and detected with an iXon Ultra 897 electron multiplied charge-coupled device (EMCCD) camera (512 × 512 pixels, Andor). For TIRFM, the penetration depth was 100 nm. The iLas2 illumination system was used for TIRFM and wide-field imaging. For experiments with RNb-Perceval-HR, a 150× oil-immersion TIRF objective (NA 1.45) and a Prime 95B Scientific metal-oxide-semiconductor (CMOS) camera (512 × 512 pixels, Photometrics) were used.

For CALI and LOV2/zdk1 experiments, the 488-nm laser in the upright position delivered an output at the objective of 2.45 mW (PM100A power meter, Thor Labs, Newton, NJ, USA). For CALI, a single flash of 488-nm laser illumination (3-s duration) was applied, with 10-ms exposures to 488-nm laser immediately before and after the CALI flash to allow imaging of SNAP-Cell-fluorescein (i.e. 3.02 s total CALI flash). For LOV2/zdk1 experiments, repeated flashes of 488-nm light (1-s duration each) were used at 2-s intervals to allow imaging with 561-nm laser illumination during the intervening periods.

Before analysis, all fluorescence images were corrected for background by subtraction of fluorescence detected from a region outside the cell. Image capture and processing used MetaMorph Microscopy Automation and Image Analysis Software (Molecular Devices) and Fiji [95]. Particle tracking used the TrackMate ImageJ plugin [96], with an estimated blob diameter of 17 pixels and a threshold of 5 pixels. Co-localization analysis used the JACoP ImageJ plugin [97]. Pearson’s correlation coefficient (*r*) was used to quantify colocalization. We report *r* values only when the Costes’ randomization-based colocalization value (P-value = 100 after 100 iterations) confirmed the significance of the original colocalization. Where example images are shown, they are representative of at least three independent experiments (individual plates of cells from different transfections and days).

### Statistics

Results are presented as mean ± SEM for particle-tracking analyses and mean ± SD for colocalization analyses, from *n* independent analyses (individual plates of cells from different transfections). Statistical comparisons used paired or unpaired Student’s *t*-tests, or analysis of variance with the Bonferroni correction used for multiple comparisons. \**p* < 0.05 was considered significant.

## Abbreviations

BFP: blue fluorescent protein
[Ca^2+^]_c_: cytosolic free Ca^2+^ concentration
CALI: chromophore-assisted light inactivation
CaM: calmodulin
CFP: cyan fluorescent protein
ER: endoplasmic reticulum
FKBP: FK506-binding protein
FRB: FKBP-rapamycin-binding domain
FP: fluorescent protein
GFP: green fluorescent protein
GNb: GFP-binding nanobody
HBS: HEPES-buffered saline
IP_3_R: inositol 1,4,5-trisphosphate receptor
LAMP1: lysosomal membrane protein 1
mCherry: monomeric Cherry
MCS: membrane contact site
MHBS: modified HBS
MP: multimerizing protein
mRFP: monomeric red fluorescent protein
OMM: outer mitochondrial membrane
PM: plasma membrane
RFP: red fluorescent protein
RNb: RFP-binding nanobody
ROI: region of interest
SOCE: store-operated Ca^2+^ entry
TIRFM: total internal reflection fluorescence microscopy
YFP: yellow fluorescent protein

## Declarations

### Availability of data and materials

All plasmids and data generated or analysed during this study are included in this published article and its supplementary information files. Plasmids are available from the corresponding authors on request and from Addgene.

### Authors’ contributions

DLP and CWT conceived the work. DLP conducted all experiments and analysis. DLP and CWT interpreted data and wrote the manuscript. DLP and CWT read and approved the final manuscript.

## Acknowledgements

Not applicable.

## Ethics Approval and Consent to Participate

Not applicable

## Additional files

### Additional file 1 (.pptx)

**Figures S1-S6:** Fig. S1 – Targeting RNb-GEMGECO Ca^2+^ sensor to RFP-tagged proteins. Fig. S2 – Targeting CALI to lysosomes with SNAP-Cell-fluorescein: cytosolic controls. Fig. S3 – Rapamycin alone does not recruit RFP-tagged or GFP-tagged proteins to mitochondria. Fig. S4 – Recruitment of proteins to native PM-mitochondria MCS using GNb-FKBP. Fig. S5 – Inducible crosslinking of RFP-tagged and GFP-tagged proteins with RNb-FKBP and GNb-FRB. Fig. S6 – DNA sequences encoding the nanobodies used.

**Fig. S1 Targeting RNb-GEMGECO Ca^2+^ sensor to RFP-tagged proteins.** (**A**) Schematic of RNb-GEMGECO fusion binding to RFP. (**B**) HeLa cells co-expressing RNb-GEMGECO and TOM20-mCh were imaged in HBS using TIRFM. Images are shown before and after addition of histamine (100 µM) and then ionomycin (5 µM). The TOM20-mCh and merged images are before additions of histamine and ionomycin. The yellow boxed region in shown enlarged in (C). Scale bar 10 μm. (**C**) Enlarged regions from (B). Scale bar 2.5 µm. (**D**, **E**) Representative timecourses of histamine and ionomycin-evoked changes in fluorescence (D) and fluorescence emission ratio (R/R_0_, where R = F_480_/F_525_) (E) of mitochondrially targeted RNb-GEMGECO. Results are representative of cells from 4 independent experiments. Relates to Figs. 3 and 4.

**Fig. S2 Targeting CALI to lysosomes with SNAP-Cell-fluorescein: cytosolic controls.** (**A**) Schematic of cytosolic SNAPf, which does not bind to RFP, after its labelling with SNAP-Cell-fluorescein. (**B-D**) HeLa cells co-expressing LAMP1-mCh and cytosolic SNAPf (Cyt-SNAPf) were treated with SNAP-Cell-fluorescein (0.5 µM, 30 min, 37°C) and imaged using TIRFM. Scale bar 10 µm. Cells were then exposed to 488-nm light for 3 s to induce CALI. Images show a representative cell at different times before (C) and after (D) CALI, with the image at t = 0 s shown in magenta and the image at t = 60 s in green. White in the temporal merged image indicates immobile lysosomes, while green and magenta indicate lysosomes that moved during the 60 s between images. Yellow boxes show regions enlarged in subsequent images. Scale bars 10 µm (main images) and 2.5 μm (enlargements). For clarity, images were auto-adjusted for brightness and contrast (ImageJ) to compensate for bleaching of mCh during tracking and CALI. (**E**) Displacements of individual lysosomes during a 60-s recording (determined by TIRFM using TrackMate, with images taken every 1 s; mean values shown by bars) for a representative HeLa cell co-expressing LAMP1-mCh and cytosolic SNAPf before and after CALI (3-s exposure to 488-nm light). Typical of n = 6 cells. Summary data are shown in Fig. 8F. Relates to Fig. 8.

**Fig. S3 Rapamycin alone does not recruit RFP-tagged or GFP-tagged proteins to mitochondria.** (**A, B**) HeLa cells co-expressing mitochondrial TOM70-mCh-FRB and mCh-Sec61β were imaged using TIRFM before (A) and after (B) addition of rapamycin (100 nM, 10 min). (**C, D**) HeLa cells co-expressing endogenously tagged GFP-IP_3_R1 and mitochondrial TOM70-mCh-FRB were imaged using TIRFM before (C) and after (D) addition of rapamycin (100 nM, 10 min). (**E**) HeLa cells co-expressing GFP-calmodulin (GFP-CaM) and mitochondrial TOM70-mCh-FRB were imaged using TIRFM before and after addition of rapamycin (100 nM, 10 min). Results are each representative of cells from 3-5 independent experiments. Scale bars 10 µm (main images) and 2.5 µm (enlargements of boxed regions). Relates to Fig. 10.

**Fig. S4 Recruitment of proteins to native PM-mitochondria MCS using GNb-FKBP.** (**A**) Schematic of GNb-FKBP fusion bound to GFP. (**B**) TIRFM images of COS-7 cells co-expressing GNb-FKBP, β_2_AR-GFP and TOM70-mCh-FRB. A representative cell (n = 3) is shown before (top row) and at the indicated times after addition of rapamycin (100 nM). Scale bar 10 µm. (**C**) Enlargements of the boxed regions in (B). Scale bar 3.75 µm. Relates to Fig. 12.

**Fig. S5 Inducible crosslinking of RFP-tagged and GFP-tagged proteins with RNb-FKBP and GNb-FRB.** (**A**) Schematic of the nanobody fusions used, with rapamycin shown as a blue sphere. (**B, C**) HeLa cells co-expressing RNb-FKBP, GNb-FRB, TOM20-GFP and mCh-Sec61β were imaged using TIRFM. A representative cell (n = 3) is shown before (B) and after (C) treatment with rapamycin (100 nM, 10 min). Scale bars 10 µm (main images) and 2.5 µm (enlargements of boxed regions). Relates to Fig. 16.

**Fig. S6. DNA sequences encoding the nanobodies used.**

**Additional file 2 (.wmv): Video 1. RNb-GGECO1.2 detects changes in** [**Ca^2+^**] **at the surface of mitochondria expressing TOM20-mCh.** The top panel shows RNb-GGECO1.2 fluorescence (488-nm TIRFM excitation) and the bottom panel shows TOM20-mCh fluorescence (561-nm TIRFM excitation). In response to histamine (100 µM, added at 60 s), local rises in [Ca^2+^]_c_ were detected at the surfaces of individual mitochondria, but not in the bulk cytosol. Ionomycin (5 µM) was added at 3 min. Video was acquired at 1 Hz and is shown at 30 frames per second (fps). Clock is in min:s. Relates to Fig. 3D.

**Additional file 3 (.wmv): Video 2. GNb-LARGECO1.2 detects local changes in** [**Ca^2+^**] **at the surface of mitochondria expressing TOM20-GFP.** The video shows GNb-LARGECO1.2 fluorescence (488-nm TIRFM excitation). Histamine (100 µM, added at 60 s) causes local rises in [Ca^2+^]_c_ at the OMM of individual mitochondria in the perinuclear region (cyan box in Fig. 4D), but not in peripheral regions (e.g. yellow box in Fig. 4D). Ionomycin (5 µM) was added at 3 min. Video was acquired at 1 Hz and is shown at 33 fps. Clock is in min:s. Relates to Fig. 4D-F.

**Additional file 4 (.wmv): Video 3. Effect of targeted CALI on lysosomal motility.** HeLa cells expressing LAMP1-mCh and RNb-SNAPf were imaged using TIRFM and 561-nm laser illumination before (top) and after (bottom) CALI (3.02 s exposure to 488-nm epifluorescence laser illumination). Video was acquired at 0.5 Hz and is shown at 3 fps. Clock is in min:s. Relates to Fig. 8.

**Additional file 5 (.wmv): Video 4. RNb-FKBP rapidly sequesters an ER integral membrane protein at the OMM.** TIRFM images of HeLa cells expressing TOM70-GFP-FRB, RNb-FKBP and mCh-Sec61β were treated with rapamycin (100 nM, added at 60 s). The ER membrane protein, mCh-Sec61β, is then rapidly sequestered at the OMM. Video was acquired at 0.5 Hz and shown at 33 fps. Clock is in min:s. Relates to Fig. 10C and D.

**Additional file 6 (.wmv): Video 5. GNb-FKBP rapidly sequesters endogenously tagged GFP-IP_3_R1 at the OMM.** TIRFM images show HeLa cells with endogenously GFP-tagged IP_3_R1 and transiently expressing TOM70-mCh-FRB and GNb-FKBP, and then treated with rapamycin (100 nM, added at 60 s). GFP-IP_3_R1 is rapidly sequestered at the OMM. Video was acquired at 0.5 Hz and is shown at 33 fps. Clock is in min:s. Relates to Fig. 10F and G.

**Additional file 7 (.wmv): Video 6. GNb-FKBP rapidly sequesters GFP-CaM at the OMM.** Epifluorescence microscopy images show HeLa cells transiently expressing GFP-CaM, GNb-FKBP and TOM20-mCh-FRB, and then treated with rapamycin (100 nM, added at 30 s). GFP-CaM is rapidly sequestered at the OMM. Video was acquired at 0.5 Hz and is shown at 9 fps. Clock is in min:s. Relates to Fig. 10H.

**Additional file 8 (.wmv): Video 7. RNb-FKBP recruits a PM protein to the OMM in response to rapamycin.** TIRFM images of HeLa cells expressing TOM70-GFP-FRB, RNb-FKBP and the PM protein, β_2_AR-mCh, and then exposed to rapamycin (100 nM, added at 60 s). There is a rapid translocation of β_2_AR-mCh to the OMM. Video was acquired at 0.5 Hz and is shown at 33 fps. Clock is in min:s. Relates to Fig. 12B-E.

**Additional file 9 (.wmv): Video 8. Crosslinking GNb-FKBP and RNb-FRB with rapamycin recruits mCh-Sec61β to TOM20-GFP in the OMM.** HeLa cells expressing GNb-FKBP, RNb-FRB, mCh-Sec61β and TOM20-GFP were stimulated with rapamycin (100 nM, added at 100 s). The TIRFM images show rapid recruitment of mCh-Sec61β to the OMM. Video was acquired at 0.2 Hz and is shown at 8 fps. Clock is in min:s. Relates to Fig. 16.

