## Additional file 1: Figures S1-S6 for "A genetically-encoded toolkit of functionalized nanobodies against fluorescent proteins for visualizing and manipulating intracellular signalling"

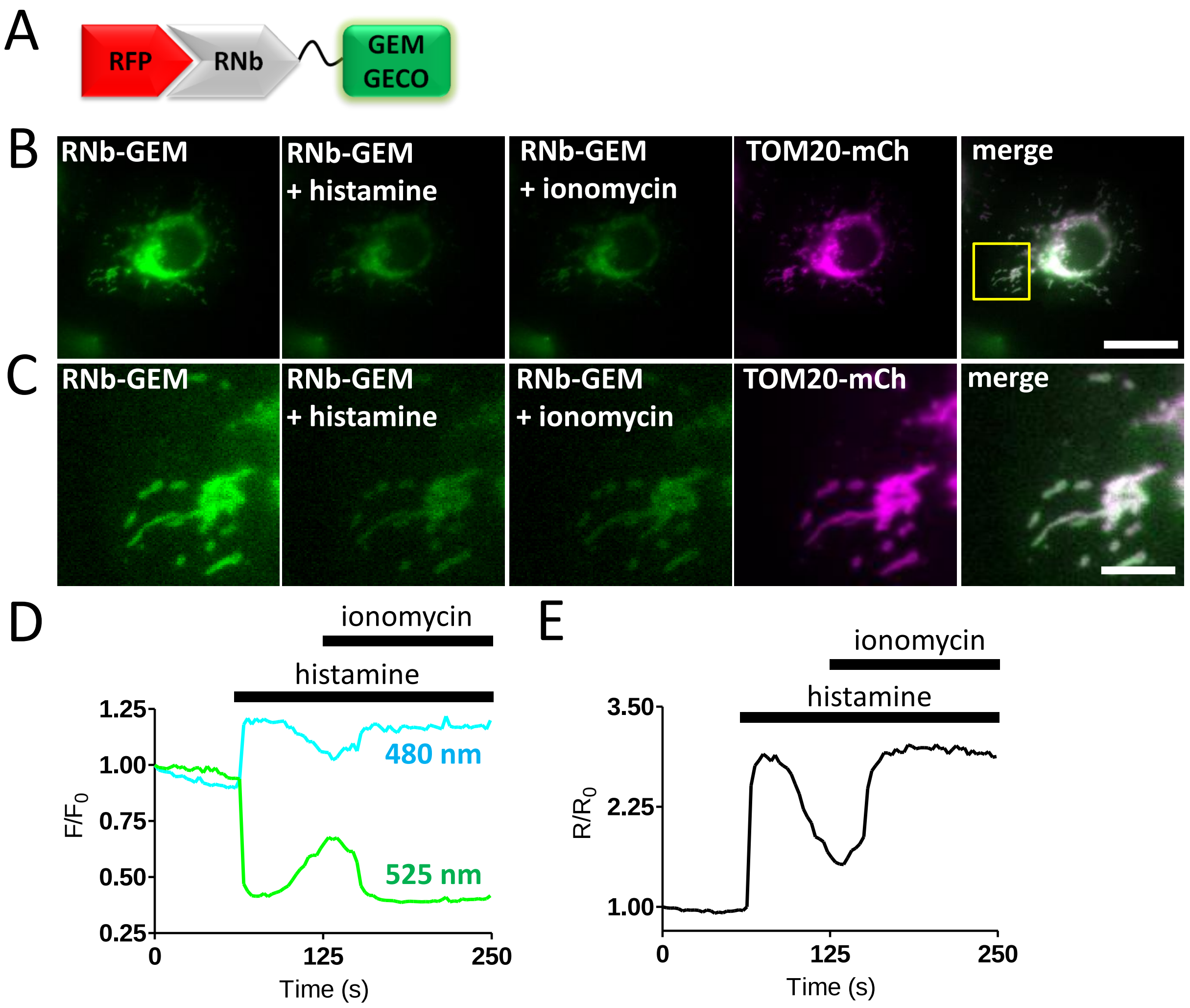


**Additional file 1: Figure S1. Targeting RNb-GEMGECO $Ca^{2+}$ sensor to RFP-tagged proteins.** (**A**) Schematic of RNb-GEMGECO fusion binding to RFP. (**B**) HeLa cells co-expressing RNb-GEMGECO and TOM20-mCh were imaged in HBS using TIRFM. Images are shown before and after addition of histamine (100 µM) and then ionomycin (5 µM). The TOM20-mCh and merged images are before additions of histamine and ionomycin. The yellow boxed region in shown enlarged in (C). Scale bar 10 µm. (**C**) Enlarged regions from (B). Scale bar 2.5 µm. (**D**, **E**) Representative timecourses of histamine and ionomycin-evoked changes in fluorescence (D) and fluorescence emission ratio ($R/R_0$, where $R = F_{480}/F_{525}$) (E) of mitochondrially targeted RNb-GEMGECO. Results are representative of cells from 4 independent experiments. Relates to ***Figs. 3*** and ***4***.

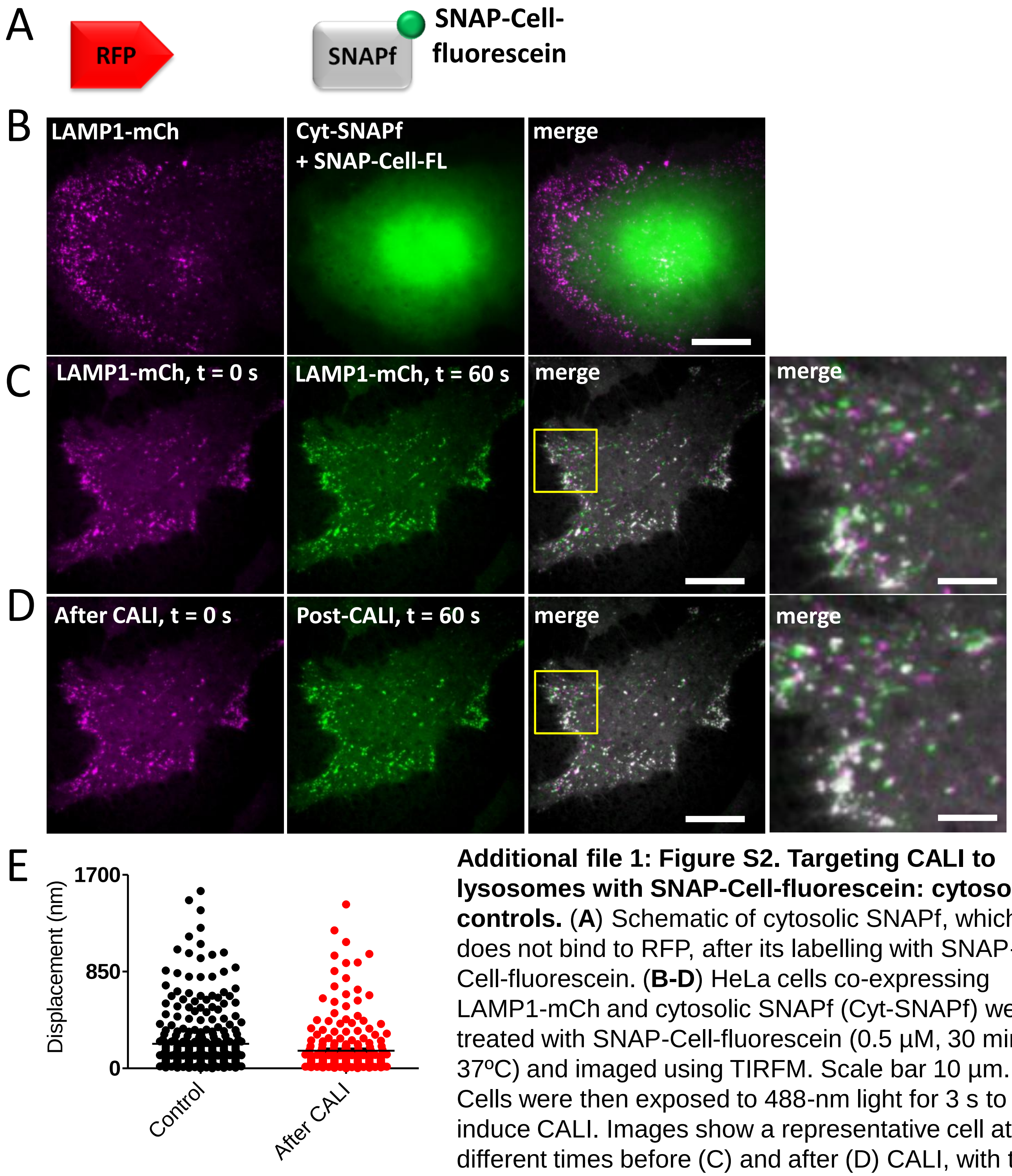


**Additional file 1: Figure S2. Targeting CALI to lysosomes with SNAP-Cell-fluorescein: cytosolic controls.** (**A**) Schematic of cytosolic SNAPf, which does not bind to RFP, after its labelling with SNAP-Cell-fluorescein. (**B-D**) HeLa cells co-expressing LAMP1-mCh and cytosolic SNAPf (Cyt-SNAPf) were treated with SNAP-Cell-fluorescein (0.5 µM, 30 min, 37ºC) and imaged using TIRFM. Scale bar 10 µm. Cells were then exposed to 488-nm light for 3 s to induce CALI. Images show a representative cell at different times before (C) and after (D) CALI, with the image at t = 0 s shown in magenta and the image at t = 60 s in green. White in the temporal merged image indicates immobile lysosomes, while green and magenta indicate lysosomes that moved during the 60 s between images. Yellow boxes show regions enlarged in subsequent images. Scale bars 10 µm (main images) and 2.5 µm (enlargements). For clarity, images were auto-adjusted for brightness and contrast (ImageJ) to compensate for bleaching of mCh during tracking and CALI. (**E**) Displacements of individual lysosomes during a 60-s recording (determined by TIRFM using TrackMate, with images taken every 1 s; mean values shown by bars) for a representative HeLa cell co-expressing LAMP1-mCh and cytosolic SNAPf before and after CALI (3-s exposure to 488-nm light). Typical of n = 6 cells. Summary data are shown in ***Fig. 8F***. Relates to ***Fig. 8***.

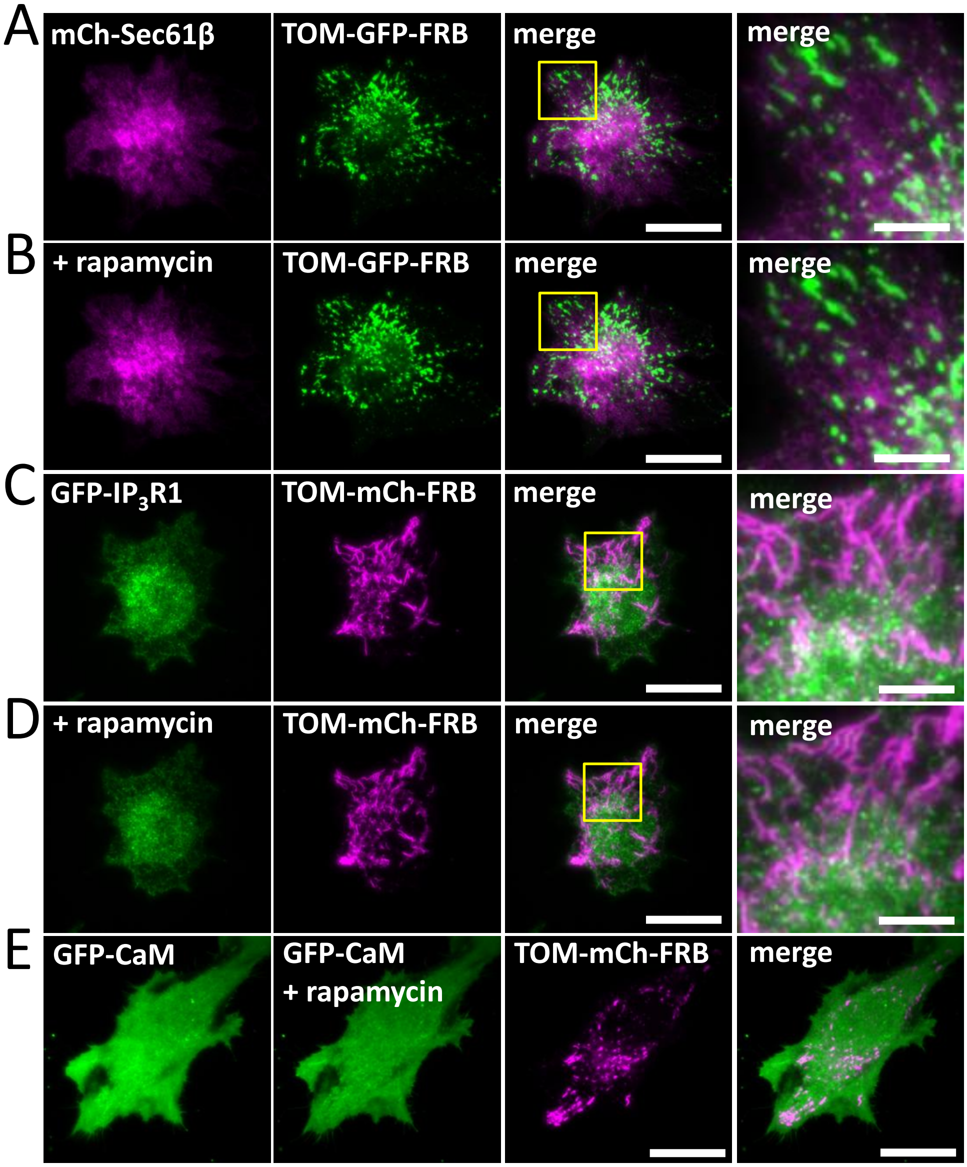


**Additional file 1: Figure S3. Rapamycin alone does not recruit RFP-tagged or GFP-tagged proteins to mitochondria.** (**A, B**) HeLa cells co-expressing mitochondrial TOM70-mCh-FRB and mCh-Sec61β were imaged using TIRFM before (A) and after (B) addition of rapamycin (100 nM, 10 min). (**C, D**) HeLa cells co-expressing endogenously tagged GFP-IP$_3$R1 and mitochondrial TOM70-mCh-FRB were imaged using TIRFM before (C) and after (D) addition of rapamycin (100 nM, 10 min). (**E**) HeLa cells co-expressing GFP-calmodulin (GFP-CaM) and mitochondrial TOM70-mCh-FRB were imaged using TIRFM before and after addition of rapamycin (100 nM, 10 min). Results are each representative of cells from 3-5 independent experiments. Scale bars 10 µm (main images) and 2.5 µm (enlargements of boxed regions). Relates to ***Fig. 10***.

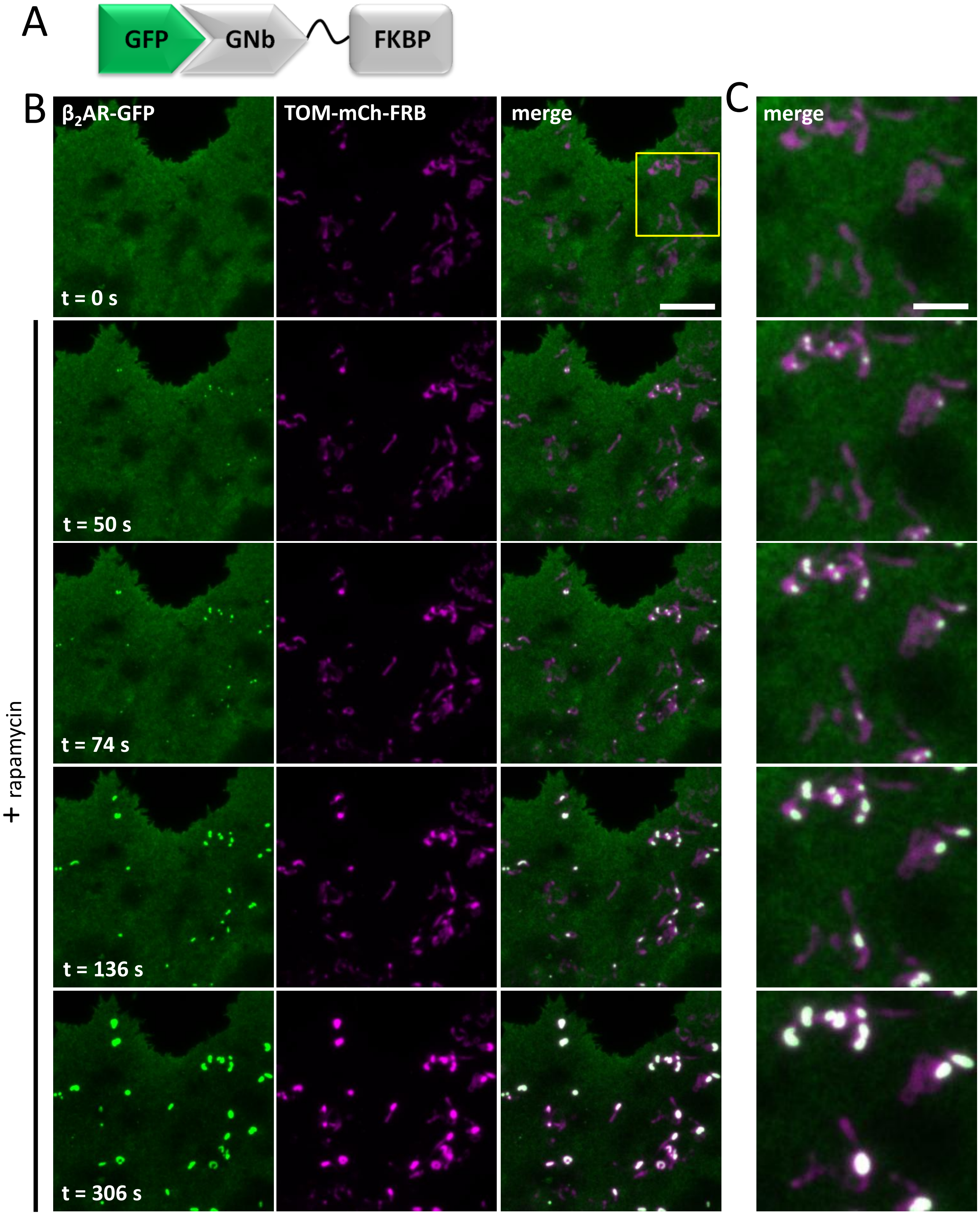


**Additional file 1: Figure S4. Recruitment of proteins to native PM-mitochondria MCS using GNb-FKBP.** (**A**) Schematic of GNb-FKBP fusion bound to GFP. (**B**) TIRFM images of COS-7 cells co-expressing GNb-FKBP, $\beta_2$AR-GFP and TOM70-mCh-FRB. A representative cell (n = 3) is shown before (top row) and at the indicated times after addition of rapamycin (100 nM). Scale bar 10 µm. (**C**) Enlargements of the boxed regions in (B). Scale bar 3.75 µm. Relates to ***Fig. 12***.

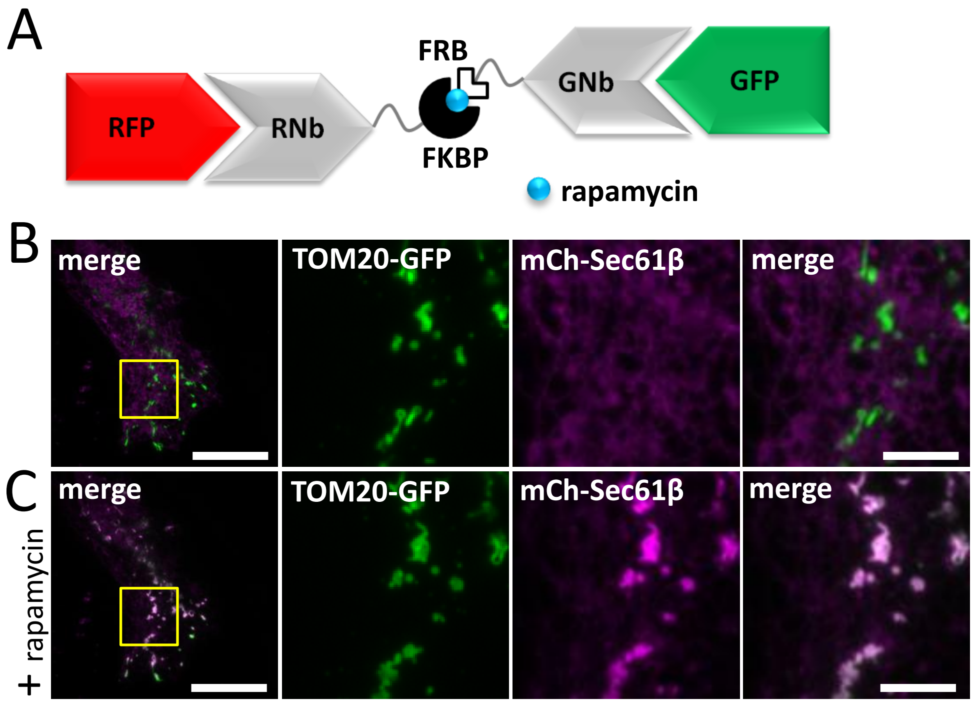


**Additional file 1: Figure S5. Inducible crosslinking of RFP-tagged and GFP-tagged proteins with RNb-FKBP and GNb-FRB.** (**A**) Schematic of the nanobody fusions used, with rapamycin shown as a blue sphere. (**B, C**) HeLa cells co-expressing RNb-FKBP, GNb-FRB, TOM20-GFP and mCh-Sec61β were imaged using TIRFM. A representative cell (n = 3) is shown before (B) and after (C) treatment with rapamycin (100 nM, 10 min). Scale bars 10 µm (main images) and 2.5 µm (enlargements of boxed regions). Relates to ***Fig. 16***.

**Additional file 1: Figure S6.** DNA sequences encoding the nanobodies used.

ATG – start codon of region encoding nanobody

**NANOBODY** – region encoding nanobody

LINKER – region encoding flexible linker between nanobody and functional module

TXX – stop codon

**GNb**

AAGCTTGCCACCATGGCTCAGGTGCAGCTGGTGGAATCTGGCGGCAGACTGGTGCAGGC
CGGCGATAGCCTGAGACTGTCTTGTGCCGCCAGCGGCAGAACCTTCAGCACATCTGCCAT
GGCCTGGTTCAGACAGGCCCCTGGCCGCGAGAGGGAATTTGTGGCCGCCATCACATGGA
CCGTGGGCAACACCATCCTGGGCGACAGCGTGAAGGGCCGGTTCACCATCAGCCGGGAC
AGAGCCAAGAACACCGTGGACCTCCAGATGGACAACCTGGAACCCGAGGACACCGCCGT
GTACTACTGCTCCGCCAGATCCAGAGGCTACGTGCTGTCCGTGCTGCGGAGCGTGGACAG
CTACGATTATTGGGGCCAGGGCACCCAAGTGACCGTGTCTGGCGGCGGAGGAAGCGGAG
GCGGAGGATCTGGGGGAGGCGGCAGTGGCGGAGGGGGATCTGGATCC

**RNb**

AAGCTTGCCACCATGGCTCAGGTGCAGCTGGTGGAAAGCGGCGGCTCTCTGGTGCAGCC
TGGCGGATCTCTGAGACTGAGCTGTGCCGCCAGCGGCAGATTTGCCGAGAGCAGCAGCA
TGGGCTGGTTCAGACAGGCCCCTGGCAAAGAACGCGAGTTCGTGGCCGCCATCTCTTGG
AGCGGCGGAGCCACCAATTACGCCGATAGCGCCAAGGGCCGGTTCACCCTGAGCCGGGA
CAACACCAAGAACACCGTGTACCTGCAGATGAACAGCCTGAAGCCCGACGACACCGCCG
TGTACTACTGCGCCGCCAACCTGGGCAACTACATCAGCAGCAACCAGCGGCTGTACGGC
TACTGGGGCCAGGGAACACAAGTGACCGTGTCCAGCCCTTTCACAGGCGGCGGAGGATC
TGGCGGAGGCGGATCTGGGGGCGGAGGCTCTGGATCC

**GNb-RNb**

AAGCTTGCCACCATGGCTCAGGTGCAGCTGGTTGAATCTGGCGGCAGACTGGTTCAGGCC
GGCGATTCTCTGAGACTGTCTTGTGCCGCCAGCGGCAGAACCTTTAGCACATCTGCCATG
GCCTGGTTCAGACAGGCCCCTGGAAGAGAACGGGAATTTGTGGCCGCCATCACCTGGAC
CGTGGGCAATACCATCCTGGGCGATAGCGTGAAGGGCAGATTCACCATCAGCCGGGACA
GAGCCAAGAACACCGTGGACCTCCAGATGGACAACCTGGAACCTGAGGACACCGCCGTG
TACTACTGCTCCGCTAGAAGCAGAGGCTACGTGCTGTCCGTGCTGAGAAGCGTGGACAG
CTACGATTATTGGGGCCAGGGCACCCAAGTGACCGTTTCTGGTGGCGGAGGATCTGGCG
GAGGTGGAAGCGGCGGAGGCGGTAGCGGAGGTGGTGGATCTGGATCCATGGCCCAGGTC
CAGCTCGTGGAAAGTGGCGGATCTCTGGTTCAACCTGGCGGAAGCCTGAGACTGAGCTG
TGCCGCTTCTGGCAGATTTGCCGAGAGCAGCAGCATGGGCTGGTTTAGGCAAGCCCCAG
GCAAAGAGAGAGAGTTCGTCGCCGCCATCTCTTGGAGTGGCGGAGCCACCAATTACGCC
GATTCTGCCAAAGGCCGGTTCACCCTGAGCAGAGACAACACAAAGAATACGGTGTATCT
CCAGATGAACTCCCTGAAGCCAGACGATACAGCCGTGTATTATTGCGCCGCCAACCTGG
GCAACTACATCAGCAGCAACCAGCGGCTGTACGGCTACTGGGGACAGGGAACACAAGTC
ACAGTGTCTAGCCCCTTCACCTGACTGCAGATATC
